# Sexually dimorphic regulation of gonadotrope cell hyperplasia in medaka pituitary via mitosis and transdifferentiation

**DOI:** 10.1101/2022.12.08.519564

**Authors:** Muhammad Rahmad Royan, Daichi Kayo, Finn-Arne Weltzien, Romain Fontaine

## Abstract

The two pituitary gonadotropins, Fsh and Lh, regulate the reproductive function in vertebrates. While many studies have investigated the regulation of gonadotropin production and release by the sex steroid feedback, its role on the regulation of gonadotrope cell number remains unclear. Using medaka as a model and an optimized protocol to restore physiological sex steroids levels following gonadectomy, we show that gonadal sex steroids not only decrease *fshb* transcript levels, but also Fsh cell number in both sexes. We then investigated the origin of the Fsh cell hyperplasia induced by gonadectomy. In both sexes, BrdU incubation shows that this is achieved via Fsh cell mitosis. *In situ* hybridization reveals that new Fsh cells also originate from transdifferentiating Tsh cells in females, but not in males. Both phenomena are inhibited by sex steroid supplementation via feeding. In males (but not females), gonadectomy (without recovery with sex steroid supplementation) also reduces *sox2* transcript levels and Sox2-immunopositive population volume, suggesting that sox2-progenitors may be recruited to produce new Fsh cells. Opposite to Fsh cells, gonadectomy decreases *lhb* levels in both sexes, and levels are not restored by sex steroid supplementation. In addition, the regulation of Lh cell number also seems to be sex dependent. Removal of gonadal sex steroids stimulates Lh cell mitosis in male (like Fsh cells), but not in females. To conclude, our study provides the first evidence on sexually dimorphic mechanisms used in the fish pituitary to remodel gonadotrope populations in response to sex steroids.

**HIGHLIGHTS:**

- Supplementing gonadectomized fish with sex steroids via feeding allows for the recovery of physiological circulating levels of sex steroids.
- Gonadal sex steroids not only regulate gonadotrope cell activity, but also gonadotrope cell number.
- Removal of gonadal sex steroids induces Fsh cell hyperplasia via mitosis of Fsh cells in both sexes, and transdifferentiation of Tsh cells into bi-hormonal Tsh/Fsh cells in females only.
- Gonadectomy also reduces the number of Sox2 progenitor cells in males (but not in females), suggesting that they may be recruited to contribute to Fsh cell hyperplasia.
- Removal of gonadal sex steroids stimulates Lh cell mitosis in males, but not in females.

## INTRODUCTION

As the master gland of endocrine systems, the pituitary regulates several essential physiological functions, including growth, homeostasis, stress, and reproduction. In particular, it plays a key role in the later function by fine-tuning gonadotropin (follicle-stimulating hormone, Fsh; luteinizing hormone, Lh) synthesis and release from gonadotrope cells to control gonadal gametogenesis and steroidogenesis (Weltzien et al., 2004). In mammals, Fsh and Lh are mostly produced by the same cells (Nakane, 1970; Pope et al., 2006), while in most teleost fishes these hormones are secreted by distinct cell types (Weltzien et al., 2014; Fontaine et al., 2022), making them an ideal model to investigate gonadotrope cell regulation and plasticity.

Gonadotropin production and release are mainly controlled by hypothalamic gonadotropin-releasing hormone (Gnrh) neurons that, in teleosts, directly innervate gonadotrope cells (Kanda, 2019). The gonadotropins are then transmitted via the bloodstream to stimulate gonadal development and maturation (Yaron and Levavi-Sivan, 2011). In addition to the upstream control from the brain, the magnitude of gonadotropin production and release is also regulated by feedback from sex steroids produced by testicular Leydig cells and ovarian granulosa and theca cells, respectively, in male and female gonads (Devlin and Nagahama, 2002). This feedback acts directly on gonadotrope cells and indirectly via the brain (e.g. Gnrh neurons) which regulates pituitary gonadotrope cells (Fontaine et al., 2020c). Upon regulation by external and internal stimuli, gonadotrope cells therefore need to be plastic to maintain proper balance of gonadotropin synthesis and secretion according to life-stage dependent fluctuating physiological demands.

Sex steroids, primarily represented by 11-ketotestosterone (11-KT) in male and estradiol (E2) in female teleosts, have been shown to affect gonadotrope cell activity and gonadotropin secretion in various teleost species. Sex steroid effects involve differences depending on species, sex, physiological status, and dose or method of administration (for review, see (Fontaine et al., 2020c)). In the Japanese medaka (*Oryzias latipes*), for example, supplementation of E2 via feeding was shown to elevate *lhb* levels in females (Kanda et al., 2011), while bathing with E2 downregulated *lhb* levels in both sexes (Fontaine et al., 2019). Sex steroid (11-KT) was also suggested to cause sexual dimorphism in medaka by suppressing *fshb* transcript levels in males but not in females (Kawabata-Sakata et al., 2020). In addition, sexual dimorphism is also found in Lh cell population volumes (Royan et al., 2021) and cell numbers (Fontaine et al., 2019) where they are higher in females. The mechanisms by which gonadotrope cell numbers are regulated by sex steroids remain poorly understood, although previous studies proposed that cell mitosis might play a role (Fontaine et al., 2019). Sex steroids thus not only regulate gonadotrope cell activity, but also remodel gonadotrope cell populations by fine-tuning the cell number.

There are three hypotheses regarding the origin of new endocrine cells which would enable pituitary endocrine cell population remodeling: mitosis, transdifferentiation, and progenitor cell differentiation. Mitosis has been demonstrated in medaka gonadotropes using sex steroid bathing (Fontaine et al., 2019; 2020a), a method shown to deliver supraphysiological levels due to bioaccumulation (Kayo et al., 2020). Thus, the effect remains controversial. Transdifferentiation, which is the conversion of one cell type to another, has never been described *in vivo* in the fish pituitary. Pituitary progenitor cells identified with Sox2 immunostaining are present in adult medaka (Fontaine et al., 2019), but their role in gonadotrope remodeling and whether they are affected by sex steroids in fish remains unknown. Therefore, we aim to investigate the role sex steroids play in these three mechanisms in the fish pituitary.

In this present study, we use medaka, a small teleost fish widely used in laboratories. It has a genetic sex determination system (Matsuda et al., 2002; Nanda et al., 2002), and offers an easy access to a wide range of genetic and imaging tools, as well as molecular techniques. These include the medaka transgenic lines in which hrGfpII and DsRed2 reporter protein synthesis are controlled respectively by the endogenous medaka *lhb* (Hildahl et al., 2012) and *fshb* (Hodne et al., 2019) promoters, and the recently developed 3D pituitary atlas which facilitates visualization of all endocrine cell populations, including gonadotropes (Royan et al., 2021). It is also a powerful model for genetic and developmental studies (Powell et al., 1996; Wittbrodt et al., 2002; Hori, 2011). Considering the variation in results obtained from different protocols of sex steroid administrations, in this study we investigated sex steroid effects on gonadotrope cell regulation using gonadectomy and a carefully optimized protocol for sex steroid supplementation that allows restoration of physiological circulating levels.

## MATERIALS AND METHODS

### Animals

Juvenile (2 months) and adult (6 months) male and female d-rR strain of wild type (WT) and double transgenic (dTg) tg(*lhb*:hrGfpII/*fshb*:DsRed2) Japanese medaka (*Oryzias latipes*) were maintained in a recirculating water system (pH 7.5; 800 μS; 28 °C) with 14L:10D photoperiod. Artemia and artificial feed (Size 200-400; Zebrafeed, Sparos) were used to feed the fish three times daily. Secondary sexual characteristics were used for sex determination (Kinoshita et al., 2009). The experiments were conducted following the Norwegian regulations on experimental animal care and welfare. Experiments with gonadectomy were approved by the Norwegian Food Safety Authority (Permit number 24305).

### Gonadectomy and sex steroid supplementation

Gonadectomy (GDX), which was shown to drastically reduce circulating sex steroids for at least 4 months (Royan et al., 2020), was performed in male and female medaka as previously described (Kanda et al., 2008). Briefly, after anesthesia using 0.02% Tricaine methanesulfonate (MS222; Sigma), fertile males and females (indicated by spawning and/or the existence of oviposited eggs in the females) were incised to access the intraperitoneal cavity. The gonads were entirely removed, and the incision was sutured using a nylon thread (10-0 USP; Crownjun). The fish were kept in recovery medium (0.9% NaCl) for three days before experiment.

Sex steroids were supplemented via feeding as demonstrated previously (Kayo et al., 2020), with some adjustments. The sex steroid-containing feed was prepared by mixing 5 mg dry feed with 10-100 ng of 11-KT or E2 (Sigma) using 96% ethanol as the vehicle and then evaporated at 40 °C overnight. Due to the short half-life of sex steroids by feeding (Kayo et al., 2020), the fish were fed five hours before and sampled two hours after the onset of the dark phase (night). Tests were performed in tanks with or without water recirculation. Supplementation with 30 ng sex steroids /day was found to closely mimic the circulating sex steroid level in medaka in a circulating system (Supp. Fig. 1) and thus was used for the rest of the study. After post-surgery recovery, each fish was fed with sex steroid-supplemented or control feed (with vehicle alone) once a day for three (fluorescence *in situ* hybridization and immunofluorescence) or five (qPCR and ELISA) days.

### BrdU treatment

Marking of mitotic gonadotrope cells was performed using BrdU incubation as previously described (Fontaine et al., 2020a). The fish were treated for 6 hours with 1 mM BrdU (Sigma) in water with 0.3% dimethyl sulfoxide (DMSO; Sigma) before sampling.

We first looked at the effect of GDX on gonadotrope cell mitosis by treating sham-operated (control) and GDX adult dTg fish with BrdU three days after recovery period (6 days post-surgery in total). In a separate experiment, we looked at the effect of sex steroids on gonadotrope cell mitosis by treating with BrdU non-steroid supplemented GDX fish (control) and GDX fish fed with sex steroids for three days following recovery period.

### Blood and tissue sampling

Blood sampling was performed as previously described (Royan et al., 2020). After euthanasia with 0.02% MS222, blood was collected from the caudal vein using a glass needle (OD 1 mm, ID 0.5 mm; GD-1; Narishige) coated with 0.05 U/μl heparin sodium (Sigma) in phosphate buffered saline (PBS). The blood was stored at −80 °C until use. For transcript level analysis, pituitaries were isolated and directly transferred into tubes containing 300 μl TRIzol (Invitrogen) and 6 zirconium oxide beads (diameter 1.4 μm; Bertin Technologies). For cell imaging analysis, the brain-pituitary complexes were fixed overnight at 4 °C in 4% paraformaldehyde (PFA; Electron Microscopy 135 Sciences) in phosphate buffered saline with Tween (PBST: PBS, 0.1%; Tween-20). After washing with PBST, tissues were serially dehydrated in 25, 50, 75, 96% ethanol at RT for 10 minutes each, followed by storage in 100% methanol at −20 °C until use.

### Quantitative polymerase chain reaction (qPCR)

qPCR, including data analysis, was performed as previously described (Burow et al., 2019; Royan et al., 2021). The pituitary tissue was homogenized. The homogenate was mixed with 120 μl chloroform and centrifugated. The pellet was resuspended with 14 μl nuclease free water. The cDNA was synthetized with 98 ng RNA using SuperScript III Reverse Transcriptase (Invitrogen) and random hexamer primers (Thermofisher Scientific). cDNA samples (diluted 1:5) were run in duplicate in a total volume of 10 μl (3 μl of cDNA, 5 μM of each forward and reverse primer; **Table 1**). The qPCR parameter was set to 10 min pre-incubation at 95 °C, 42 cycles of 95 °C for 10 s, 60 °C for 10 s, and 72 °C for 15 s, followed by melting curve analysis to evaluate PCR product specificity. The relative mRNA level was calculated with *rpl7* and *gapdh* as the reference genes.

**Table 1.**
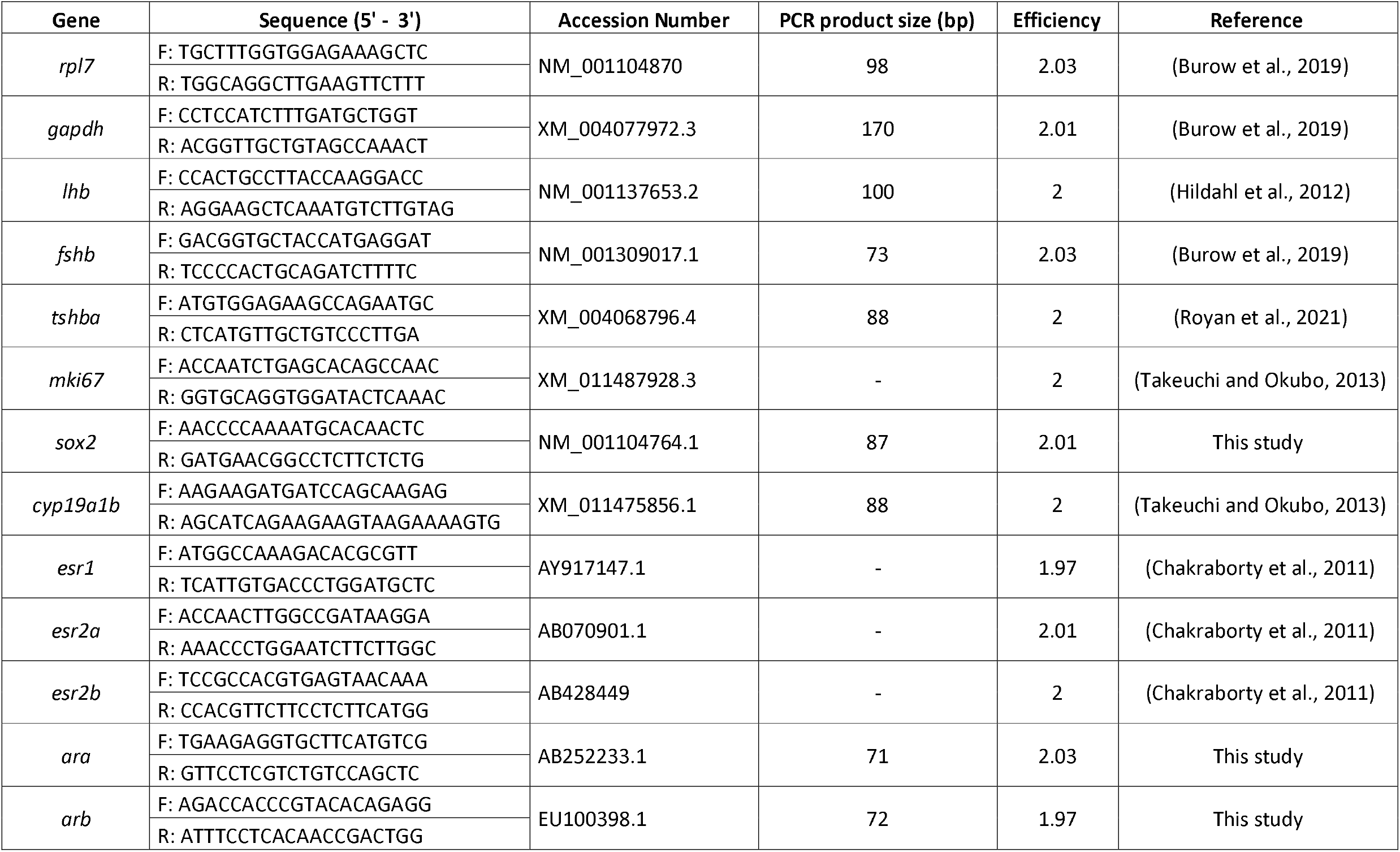
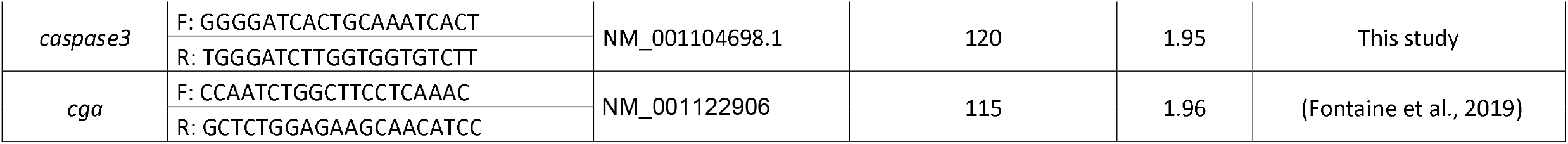
Primer sequences used for mRNA level analysis in the medaka pituitary

### Double-color fluorescence *in situ* hybridization (FISH)

FISH was performed as previously described (Fontaine et al., 2013; Royan et al., 2021) with some minor adjustments. After rehydration in a series of decreasing ethanol concentrations, the brain-pituitary complex was hybridized with previously validated specific probes (0.11 – 3.17 ng/μl) (**Table 3)** for 18 hours at 55°C and incubated with different combinations of anti-fluorescein isothiocyanate (FITC)- and anti-digoxigenin (DIG)-conjugated antibodies (Roche Diagnostics), followed by tetramethylrhodamine (TAMRA)-(Thermofisher) or FITC-conjugated tyramides (Sigma). After staining nuclei with DAPI (1:1000, 4’, 6-diamidino-2-phenylindole dihydrochloride; Sigma) and several washes with PBST, the tissue was embedded in 3% agarose (in H_2_O) and sectioned parasagitally with 70 μm thickness using a vibratome (Leica). The slices were mounted using Vectashield H-1000 Mounting Medium (Vector, Eurobio/Abcys) between microscope slides and cover slips (Menzel Glässer, VWR). For whole pituitary mounting, spacers (Reinforcement rings, Herma) were added in between.

### Immunofluorescence

Immunofluorescence was done as previously described (Fontaine et al., 2013). Following serial rehydration, the brain-pituitary complex was washed with PBST three times, embedded in 3% agarose, and para-sagittally sectioned at 70 μm thickness using a vibratome. Following an epitope retrieval treatment with 2M HCl for 1 hour at 37 °C, the free-floating sections were incubated for 1 hour at room temperature (RT) in blocking solution (4% normal goat serum, 0.5% Triton, 1% DMSO), then incubated overnight at 4 °C with anti-BrdU antibody (Abcam) and anti-medaka-Fshβ (Burow et al., 2019). For goat anti-human Sox2 (1:500; Immune Systems), the blocking solution was 0.1% Triton, 1% bovine serum albumin, and 10% normal donkey serum without pre-treatment. All antibodies used in this study have previously been validated (**Table 2**).

**Table 2.** Primary and secondary antibodies used for immunofluorescence

| Antibody | Dilution | Source | Reference |
| --- | --- | --- | --- |
| Rat anti-BrdU | 1 : 250 | abcam (ab6326) | (Fontaine et al., 2019) |
| Rabbit anti-medakaFshβ | 1 : 500 | custom-made | (Burow et al., 2019) |
| Goat anti-Sox2 | 1 : 500 | Immune systems (GT15098) | (Fontaine et al., 2019) |
| Goat anti-rabbit (Alexa-555) | 1 : 500 | Invitrogen (A21429) | (Hodne et al., 2019) |
| Donkey anti-goat (Alexa-555) | 1 : 500 | Invitrogen (A21432) | (Fontaine et al., 2019) |
| Donkey anti-rat (Alexa-647) | 1 : 500 | Invitrogen (A78947) | (Fontaine et al., 2019) |

**Table 3.**
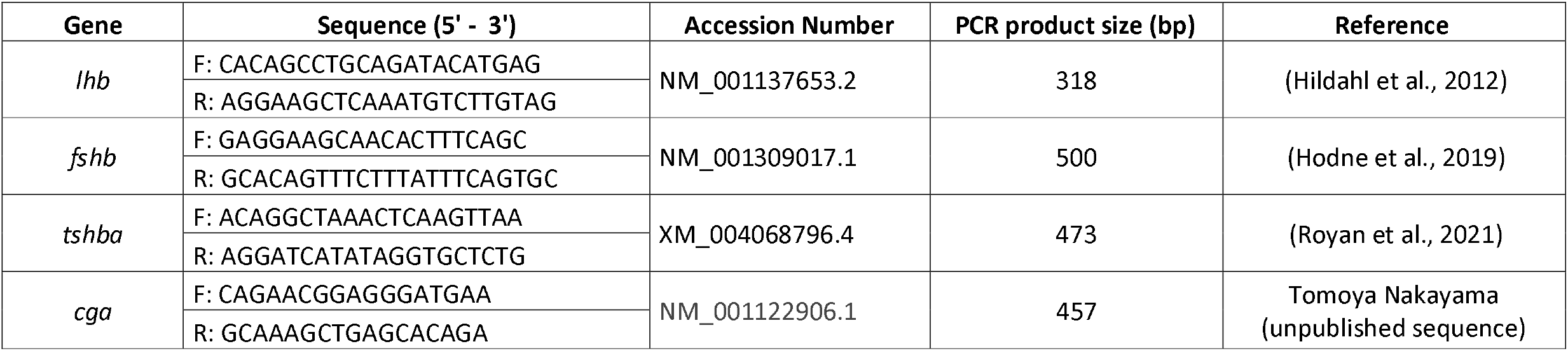
Primer sequences used to make fluorescence *in situ* hybridization (FISH) probes

### Sex steroid extraction and enzyme-linked immunosorbent assays (ELISA)

Sex steroid extraction and ELISA were performed as previously described (Kayo et al., 2020). Briefly, the blood samples were diluted 200× in PBS, extracted with diethyl ether (Sigma), and assayed using E2 and 11-KT ELISA kits (Cayman Chemical) according to manufacturer’s instructions. Sex steroid concentrations were calculated with a standard curve fitted by a 4-parameter-logistic regression (R^2^ > 0.99).

### Image acquisition, cell counting, and cell volume

Fluorescent images were generated using a Leica Confocal Microscope (DMi8, Leica) with 20× apochromat objective (numerical aperture 0.75), using lasers with wavelengths of 405 (DAPI; Alexa-405), 488 (FITC; Alexa-488), 555 (TAMRA; Alexa-555), and 647 (Red fluorophore cyanine 5, Cy5; Alexa-647). To avoid cross talk between fluorophores, the channels were obtained sequentially. Las X (v3.7, Leica) and ImageJ (1.53t; http://rsbweb.nih.gov/ij/) were used to process the images. Cell counting was performed according to (Fontaine et al., 2019) using Cell Profiler software (v2.1.0) while the double-labelled cells were manually counted using cell-counter plugin in ImageJ. Cell volume was analyzed as previously described (Royan et al., 2021), in which the absolute volume was calculated based on the area of the labeling and the voxel depth, while relative volume also includes the DAPI staining.

### Statistical analysis

Data were checked for normality using Shapiro-Wilk normality test and for homogeneity using Levene’s test. Statistical significance was set to *p* < 0.05. Differences between groups were evaluated using one-way ANOVA followed by Tukey *post hoc* test and independent-sample Student’s t-test, or Games-Howell *post hoc* test and Mann-Whitney U test for non-parametric analyses. All statistical analyses were performed using Jamovi (Version 2.2.5) (Jamovi, 2021) and graphs are provided as mean ± standard error of the mean (SEM).

## RESULTS

### Pubertal changes in gonadotrope cell activity and number, and in plasma sex steroid levels

We started by comparing the pituitary mRNA levels of *fshb* and *lhb* between juvenile and adult fish using qPCR, Lh and Fsh cell numbers using whole-mount pituitary imaging from dTg fish, and circulating sex steroid levels using ELISA. Both *fshb* and *lhb* mRNA levels (Fig. 1A-B) and Fsh and Lh cell numbers (Fig. 1C-D) increased between juvenile and adult stages. We also observed sexual dimorphism of both *fshb* (*p* < 0.001) and *lhb* (*p* = 0.002) mRNA levels and Lh cell numbers (*p* < 0.001), which are higher in females. The increase in both mRNA level and cell number from juveniles to adults coincided with an increase of 11-KT and E2 levels in males and females, respectively (Fig. 1E-F).

**Figure 1.**
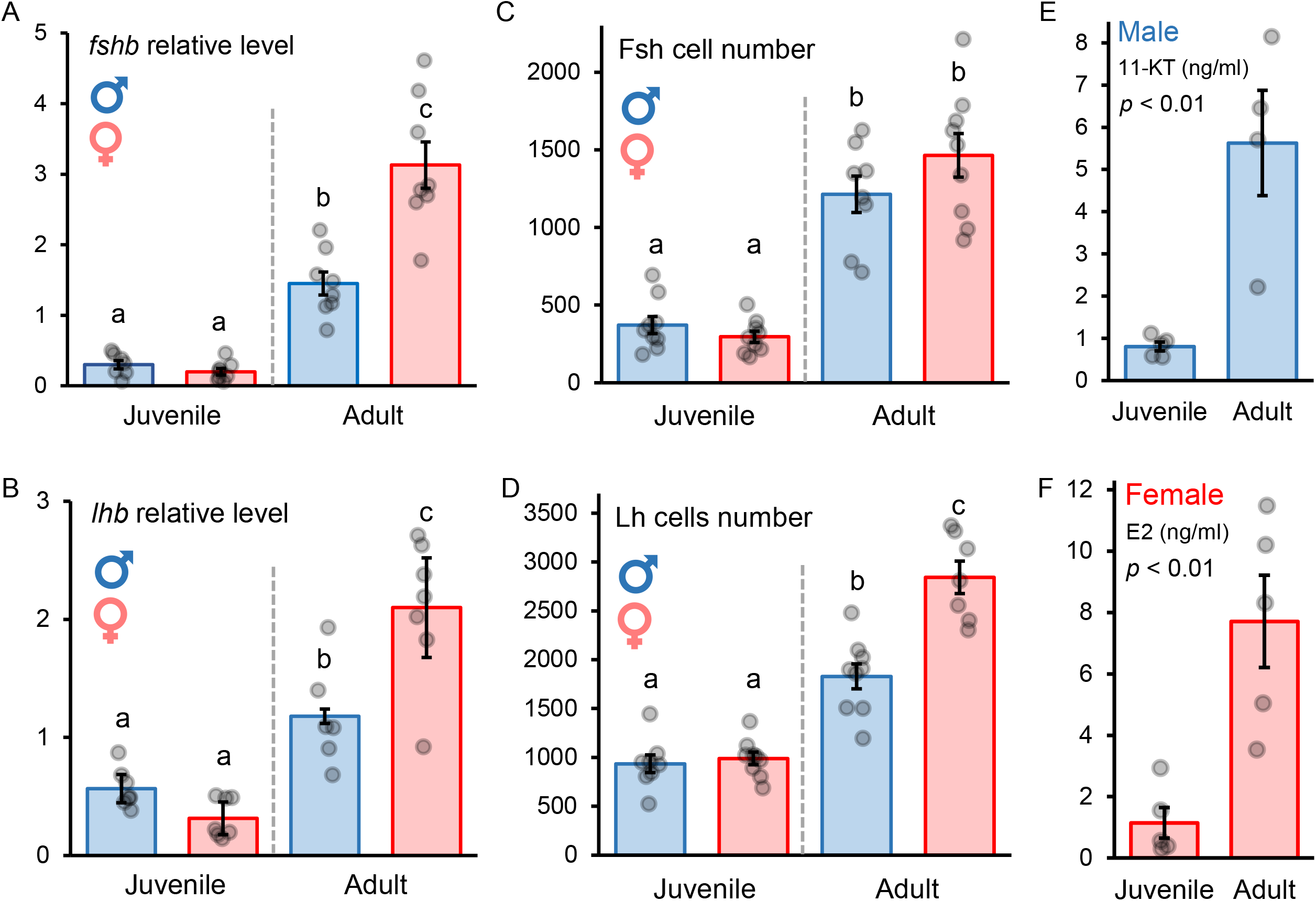
Comparison of gonadotropin mRNA levels, gonadotrope cell number, and sex steroid levels in juvenile and adult medaka. **(A-B)** Relative mRNA levels of *fshb* and *lhb* in WT medaka (n = 6 −8), and **(C-D)** Fsh and Lh cell number in double transgenic (dTg) medaka pituitary (n = 7 - 9). Different letters display statistically significant differences (*p* < 0.05) between groups as evaluated by One-way ANOVA followed by Tukey *post hoc* test. **(E-F)** Sex steroid (11-KT in males and E2 in females) levels in juvenile and adult medaka as tested using whole-blood ELISA and statistically analyzed by independent sample student’s t-test (n = 4 - 5). Graphs are provided as mean ± SEM, with jitter dots as the data points.

### Optimization of sex steroid supplementation protocol

11-KT and E2 levels showed a circadian rhythm in adult males and females respectively (Fig. 2A-B). 11-KT levels in the males gradually increased during late afternoon and evening, and peaked at around 12 am midnight (14.4 ± 1.4 ng/ml). Then, levels fell within 4 hours until the onset of the light phase (day) on at 8 am (4.9 ± 0.6 ng/ml). Meanwhile, E2 levels in females peaked between 12 am and 4 am (11.9 ± 0.7 ng/ml) and decreased to their lowest levels at 8 am in the morning (4.3 ± 0.4 ng/ml). Interestingly, in females a second peak occurred around 4 pm. Considering the timing of peak circulating sex steroids, we sampled all fish in this study two hours after the light onset of the dark phase (night) when levels are high for both 11-KT and E2.

**Figure 2.**
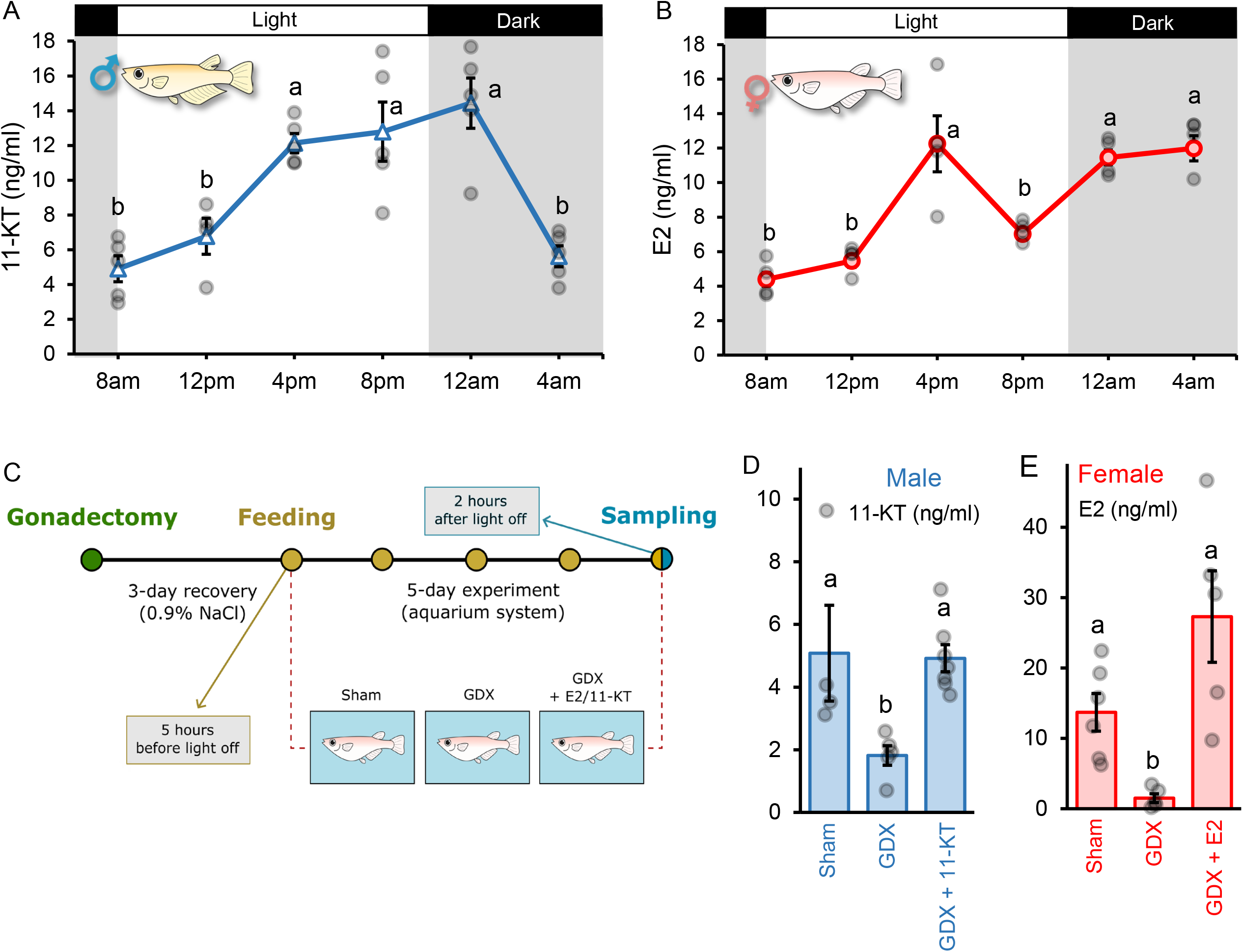
Optimization of sex steroid supplementation protocol shows the recovery of physiological level of circulating sex steroids. **(A-B)** Circadian rhythms of circulating 11-KT and E2 levels in adult medaka male and females, respectively (n = 4 - 5). **(C)** Experimental scheme illustrating gonadectomy, the duration of recovery period and feeding experiment, as well as the feeding and sampling time in the three groups of fish: sham-operated fish fed with control feed, GDX fish fed with control feed, and GDX fish fed with sex steroid supplemented feed. **(D-E)** Circulating 11-KT and E2 levels in male and female medaka, respectively, between sham-operated (Sham), gonadectomized (GDX), and fish fed with sex steroids supplemented feed (GDX + 11-KT or GDX + E2) (n = 4 - 6). Graphs are provided as mean ± SEM, with jitter dots as the data points. In all graphs, different letters display statistically significant differences (*p* < 0.05) between groups as evaluated by One-way ANOVA followed by Tukey *post hoc* test (except for E2, analyzed using Games-Howell *post hoc* test).

While gonadectomy (GDX) decreased E2 levels in females, we found that feeding with 10-30 ng of E2 while keeping the fish in a water recirculating system restored the levels of E2 observed in the sham-operated (control) fish (Supp. Fig. 1). In contrast, removal of water circulation in the tank significantly increased the circulating E2 levels measured in the blood (*p*_c-d_ < 0.001; Supp. Fig. 1). We thus supplemented GDX fish in this study with 30 ng E2/day and fed the fish five hours before the onset of the dark phase to account for the half-life of sex steroids via feeding (Kayo et al., 2020). Similar feeding protocol was used for 11-KT as this was also found to enable recovery of 11-KT circulating levels. Indeed, using this optimized feeding protocol, we observed decreased sex steroid levels in GDX fish compared to sham group (11-KT, *p* = 0.034; E2, *p* = 0.012), and recovery circulating levels of sex steroids after supplementation with 30 ng/day of 11-KT or E2 for five days (11-KT, *p* = 0.021; E2, *p* = 0.035; Fig. 2D-E).

### Sex steroids regulate gonadotrope cell activity and number

In both sexes, *fshb* mRNA levels were higher in the GDX group compared to controls (male, *p* = 0.006; female, *p* < 0.001), while sex steroid supplementation recovered *fshb* levels back to basal levels (male, *p* = 0.003; female, *p* < 0.001; Fig. 3A).

**Figure 3.**
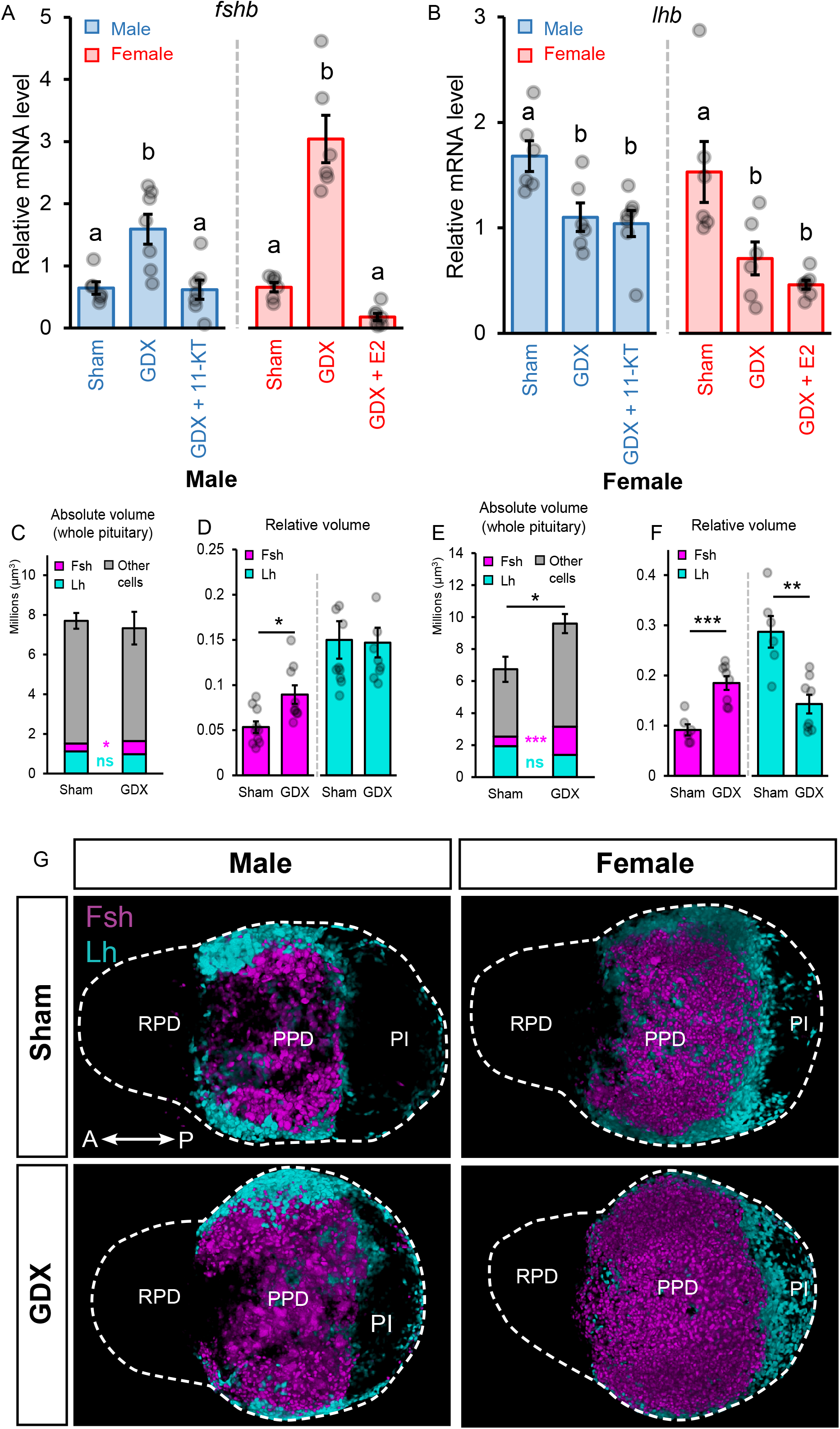
Sex steroids regulate gonadotrope cell activity and number in gonadectomized medaka. **(A-B)** Pituitary *fshb* and *lhb* levels in sham-operated (Sham), gonadectomized (GDX), and fish fed with sex steroids supplemented feed (GDX + 11-KT or GDX + E2) (n = 6 – 7). The aforementioned data were evaluated with One-way ANOVA followed by Tukey *post hoc* test, in which different letters display statistically significant differences (*p* < 0.05) between groups. **(C-F)** Absolute and relative Fsh and Lh cell volume in male **(C-D)** and female **(E-F)** double transgenic (dTg, *lhb*:hrGfpII/*fshb*:DsRed2) medaka pituitary of sham-operated (Sham) and gonadectomized (GDX) fish calculated from images taken two months after surgery (n = 8 – 10). Graphs are provided as mean ± SEM, with jitter dots as the data points. The statistical analysis was performed using independent sample student’s t-test (female) and Mann-Whitney U test (male). **(G)** 3D projections of dTg adult fish pituitary showing Fsh cells are in magenta, and Lh cells are in cyan. The pituitary is delimited with a white dashed line. Two direction arrows display the direction of the pituitary (A, anterior; P, posterior). Abbrevations: RPD, *rostral pars distalis*; PPD, *proximal pars distalis*; PI, *pars intermedia*.

Meanwhile, *lhb* mRNA levels were lower in the GDX group (male, *p* = 0.023; female, *p* = 0.017), but were not recovered by sex steroid supplementation (Fig. 3B). We thus tried different protocols previously reported to increase *lhb* levels to determine whether we could recover the GDX-induced reduction of *lhb* levels. First, an intraperitoneal injection of 15-30 μl/g fish with 1 μM Gnrh1 (previously shown to stimulate Lh cells *in vitro* (Hodne et al., 2019)) with or without 30 ng E2 also did not recover *lhb* levels in female medaka (*p* < 0.01) (Supp. Fig. 2). Second, as T was found to stimulate *lhb* levels in channel catfish (Kazeto and Trant, 2005), we used the optimized feeding protocol as described above with testosterone (T) that enables recovery of circulating T levels in GDX males. However, we found that T further decreased *lhb* levels in GDX medaka males (*p* < 0.001) (Supp. Fig. 3).

Furthermore, we found that steroid supplementation with 11-KT in males and E2 in females failed to restore transcript levels of other genes, including aromatase (*cyp19a1b*) and sex steroid receptors (*esr1, esr2a, esr2b, ara, arb*) (Supp. Fig. 4A-F). While aromatase, *ara*, and *arb* levels were rescued with E2 supplementation in females, no transcript levels recovered with 11-KT supplementation in males. While *esr1* mRNA levels decreased following GDX and did not recover with 11-KT feeding in males, they were not affected by GDX in females, and E2 increased levels compared to control. Meanwhile, neither GDX nor sex steroid supplementation affected *esr2a* or *esr2b* transcript levels.

In addition to the changes in cell activity, we observed changes in gonadotrope cell populations 2 months after GDX. Despite no detectable difference in individual cell volume (data not shown), absolute and relative Fsh cell population volumes increased in GDX males and females (male, *p-*_absolute_ = 0.043, *p-*_relative_ = 0.017; female, *p-*_absolute_ = 0.012, *p-*_relative_ < 0.001; Fig. 3C-F). However, we found that the relative Lh cell population volume significantly decreased in females (*p* = 0.001; Fig. 3F), despite no difference in the apoptotic cell marker transcript levels, *caspase3* (Supp. Fig. 4G). In males, GDX had no effect on Lh population volume. The changes in gonadotrope cell populations can be clearly visualized in 3D pituitary views, especially for Fsh cells in which more cells can easily be observed in GDX fish (Fig. 3G).

### Gonadal sex steroids inhibit gonadotrope cell mitosis

We looked at the origin of newborn Fsh cells and investigated whether they originate from cell mitosis and whether sex steroids play a regulatory role. First, we observed that the transcript levels of a cell proliferation marker gene, *mki67*, increased in GDX fish compared to sham-operated fish (male, *p* = 0.021; female, *p* = 0.008). E2 supplementation normalized *mki67* levels in females (*p* = 0.019; Fig. 4A). However, in males, 11-KT only seemed to decrease *mki67* levels. Indeed, *mki67* levels in 11-KT fed GDX fish were not significantly different from levels in control (*p* = 0.771) nor in GDX fish (*p* = 0.072). The increase of proliferative marker transcripts in GDX fish is consistent with the higher number of mitotic cells observed in the pituitary, while sex steroids suppress mitosis (Fig. 4C). In particular, Fsh cell mitosis increased in GDX fish, in both sexes (male, *p* = 0.029; female, *p* = 0.036), and decreased with sex steroid supplementation (male, *p* = 0.004; female, *p* = 0.004; Fig. 4D-E). By contrast, Lh cell mitosis significantly increased in GDX males (*p* < 0.001; Fig. 4D) and seemed to decrease with 11-KT supplementation without being significantly different from the GDX fish (*p* = 0.094). In females, no change in Lh cell mitosis was observed.

**Figure 4.**
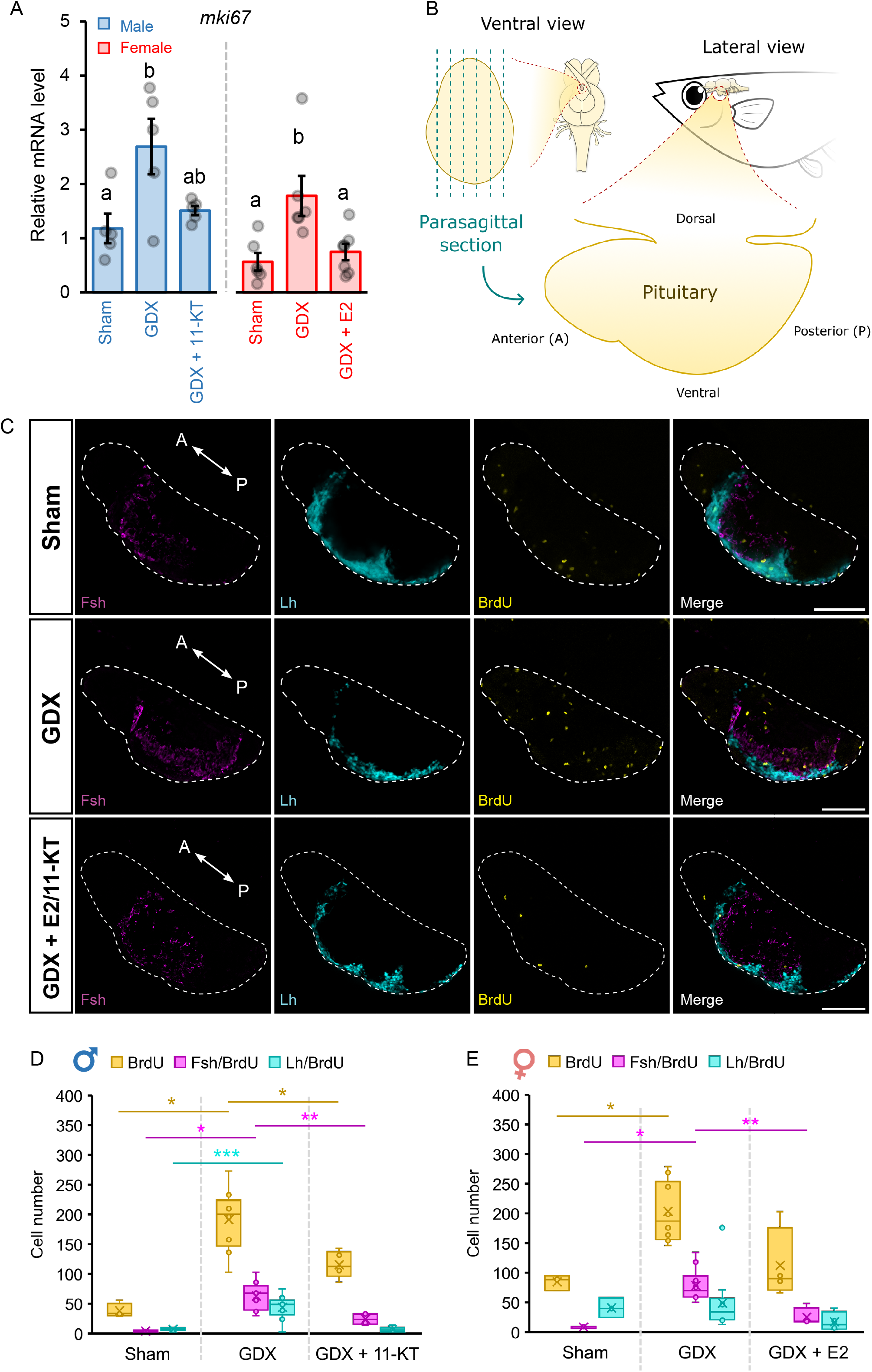
Sex steroids inhibit gonadotrope cell mitosis in medaka pituitary. **(A)** mRNA levels of *mki67*, a mitotic cell marker, in sham-operated (Sham), gonadectomized (GDX), and fish fed with sex steroids supplemented feed (GDX + 11-KT or GDX + E2). The data was analyzed by One-way ANOVA followed by Tukey *post hoc* test (n = 5 – 7). The graph is provided as mean ± SEM, with jitter dots as the data points. **(B)** A schematic illustration of the section position (parasagittal section) used for the following images. **(C)** Representative confocal planes of fish pituitary sections from sham-operated, GDX, and GDX fish fed with 11-KT or E2 supplemented feed. Tissue sections were labeled for BrdU (yellow labeling) and Fshβ by immunofluorescence (magenta). Double transgenic (dTg, *lhb*:hrGfpII/*fshb*:DsRed2) fish were used as they allow the visualization of Lh cells through Gfp (Cyan). The pituitary is delimited with a dashed line. Scale bars are in 100 μm. Two direction arrows display the orientation of the pituitary (A, anterior; P, posterior). **(D-E)** Number of cells labeled for BrdU only or together with Gfp (*lhb*) or Fshβ in sham-operated, GDX, and GDX fish fed with 11-KT or E2 supplemented feed. Significant differences were tested using Mann-Whitney U test (BrdU and Fsh) and independent sample student’s t-test (Lh/BrdU in male only) (n = 3 - 5). The graphs are provided as box and whisker plots, with the horizontal line as median, cross (×) as the mean, and dots as the data points (* < 0.05; ** < 0.01; *** < 0.001).

### Gonadal sex steroids inhibit transdifferentiation of Tsh cells into Fsh cells in females only

We then investigated whether new Fsh cells could originate from transdifferentiation of other cell types, especially those sharing the alpha subunits: Lh and Tsh cells.

Using double-color FISH for *fshb* and *lhb*, we did not observe any difference in the number of cells co-expressing *fshb* and *lhb* between control and GDX fish in both sexes after three days (Supp. Fig. 5).

*tshba*-expressing cells constitute a large cell population in the female pituitary while in males they formed a relatively small cluster (Fig. 5A). In females, *tshba* mRNA levels significantly decreased following GDX (*p* = 0.033) but did not recover with E2 supplementation (Fig. 5B). In contrast, *tshba* levels were not affected by GDX or 11-KT-supplemented feed in males.

**Figure 5.**
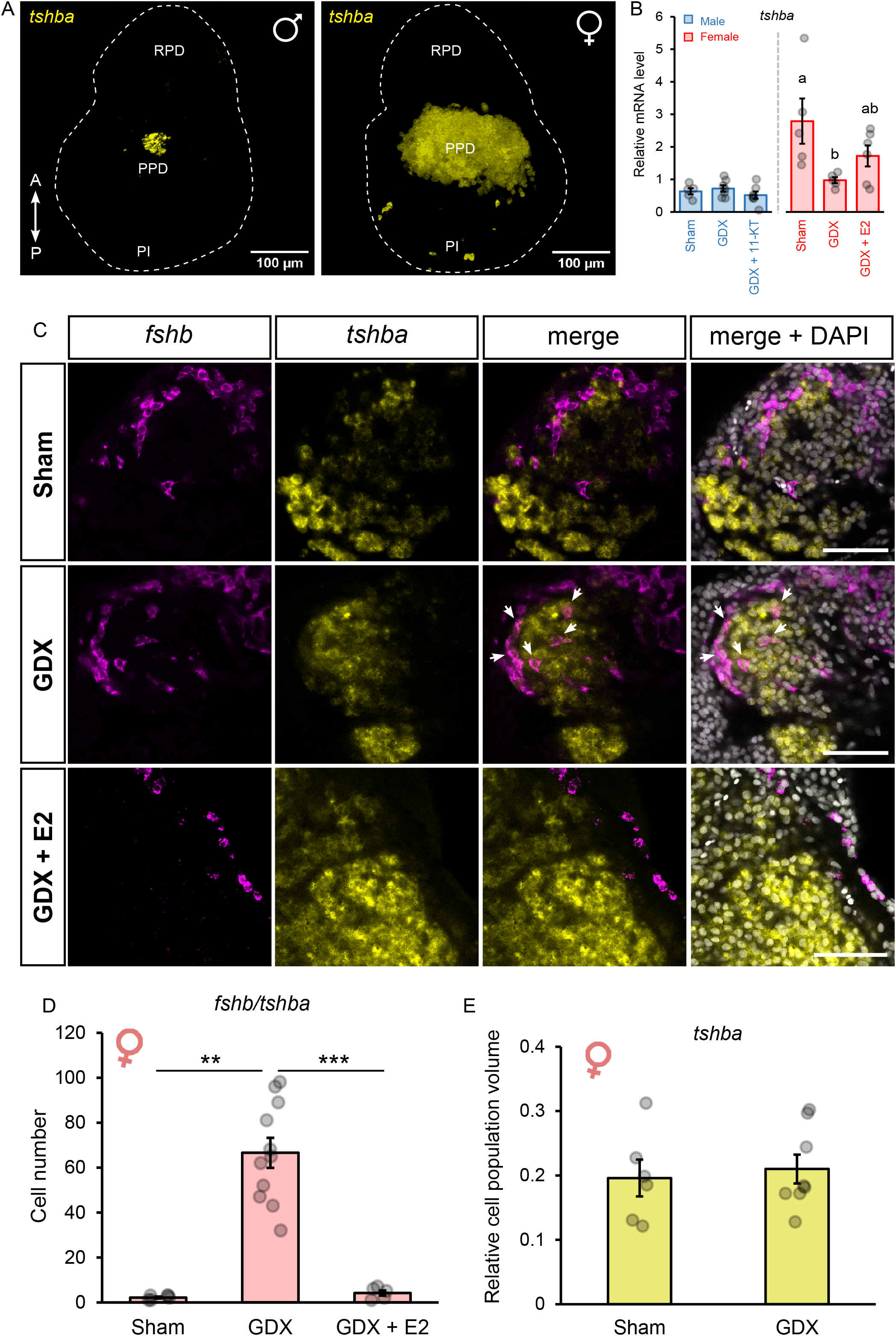
Sex steroid inhibits Tsh cell transdifferentiation to Fsh cells. **(A)** 3D reconstruction of a z-stack confocal of *tshba*-expressing cells labeled by fluorescence *in situ* hybridization in male and female medaka pituitary. **(B)** *tshba* mRNA levels in sham-operated (Sham), gonadectomized (GDX), and fish fed with E2 supplemented feed (GDX + E2) (n = 5 – 7). Statistically significant differences were evaluated by One-way ANOVA followed by Tukey *post hoc* test and displayed with different letters (*p* < 0.05) between groups. **(C)** Representative confocal planes of parasagittal sections from sham, GDX, and GDX + E2 in female fish pituitaries showing cells expressing *fshb* (magenta), *tshba* (yellow), or both together (shown by the arrows). Nuclei (grey) were labeled with DAPI staining. Abbreviations: RPD, *rostral pars distalis*; PPD, *proximal pars distalis*; PI, *pars intermedia*. Two-direction arrows display the orientation of the pituitary (A, anterior; P, posterior). Scale bars are in 50 μm. **(D)** Number of cells co-expressing *fshb* and *tshba* in sham, GDX, and GDX + E2 female fish pituitaries (n = 5 - 8). Statistically significant differences were tested by independent sample student’s t-test. **(E)** Graph showing *tshba*-expressing cell population volume in sham and GDX females six weeks post-GDX (n = 6 - 8), with statistically significant differences tested by independent sample student’s t-test. Graphs are provided as mean ± SEM, with jitter dots as the data points. (** < 0.01; *** < 0.001).

Double-color FISH for *fshb* and *tshba* revealed numerous cells labeled for both *fshb* and *tshba* in the pituitary of GDX females (Fig. 5C), something which was very rare in control and in E2-supplemented females (Fig. 5D). In males, this phenotype was rarely observed regardless of treatment (Supp. Fig. 6). Interestingly, six weeks after GDX, there was no change in *tshba*-expressing cell population volume in females (Fig. 5E), and cells expressing both *tshba* and *fshb* could still be observed (data not shown). In addition, we did not observe cells co-expressing *tshba* and *lhb* in either sex, in any condition (Supp. Fig. 7).

### Gonadectomy affects Sox2 progenitor cell number in males

Finally, we investigated whether progenitor cells could participate in Fsh proliferation following GDX. *sox2* transcript levels (marker of progenitor cells) decreased in the pituitary following GDX in males (*p* = 0.014; Fig. 6A), consistent with decreased Sox2-immunopositive cell population volume (*p* = 0.015; Fig. 6B), as seen in Fig. 6C. However, *sox2* levels did not recover with sex steroid supplementation. Meanwhile, in females, *sox2* transcript levels did not differ between control and GDX groups (Fig. 6A), consistent with the relatively stable Sox2-immunopositive cell population volume (Fig. 6B).

**Figure 6.**
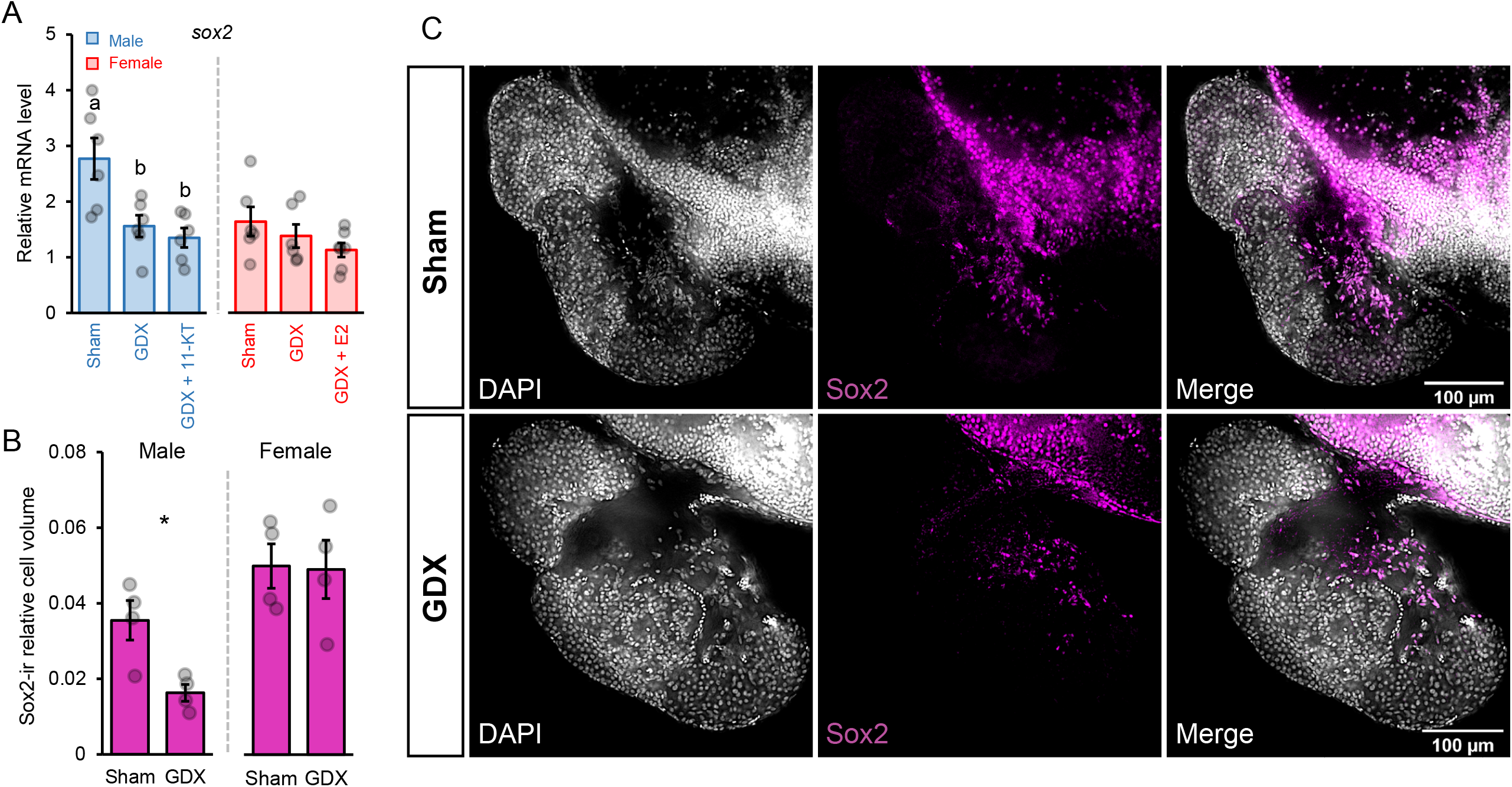
Progenitor cell population are affected by gonadectomy in males. **(A)** *sox2* mRNA levels in sham-operated (Sham), gonadectomized (GDX), and fish fed with sex steroids supplemented feed (GDX + 11-KT or GDX + E2) (n = 6 – 7). Statistically significant differences were evaluated by One-way ANOVA followed by Tukey *post hoc* test. Different letters display statistically significant differences (*p* < 0.05) between groups. Graphs are provided as mean ± SEM, with jitter dots as the data points. **(B)** Sox2 cell population volume in sham and GDX fish (n = 4). Statistically significant differences were tested with independent sample student’s t-test. **(C)** Images taken with Thunder microscope showing Sox2-immunopositive cells (magenta) in the sham and GDX male pituitary. Cell nuclei (grey) were labeled with DAPI. Scale bars: 100 μm.

The mRNA levels of *cga* (glycoprotein alpha subunit, another early marker of gonadotrope differentiation) increased in females following GDX and were normalized by E2. However, using double-color *in situ* to colocalize mixed *tshba*/*fshb*/*lhb* and *cga*, no cells expressed only *cga* in either control or GDX groups (Supp. Fig. 8A-B). Nonetheless, in GDX fish, *cga* labeling always appears brighter in the area where Tsh cells are expected (Supp. Fig. 8B).

## DISCUSSION

In teleosts, many studies have investigated the effects of sex steroids on pituitary gonadotropin production, either at the transcript level (Sohn et al., 1998; Klenke and Zohar, 2003) or at the protein level (Saligaut et al., 1998; Vetillard et al., 2003). However, little is known about how gonadotrope cell numbers are regulated. In this study, we investigated the role of sex steroids in gonadotrope cell population remodeling.

We first show the link between gonadotrope cells and sex steroids by investigating gonadotrope cell activity (transcript levels), cell number, and sex steroid levels, between juvenile and adult medaka. The higher transcript levels of *fshb* and *lhb* are in line with higher levels of 11-KT and E2 in adults compared to juveniles. This is not surprising since it is already known that Fsh and Lh play a crucial role in gametogenesis and steroidogenesis in the gonads, which is necessary for adult fish to reproduce (Takahashi et al., 2016; Kanda, 2019). The simultaneous increase of gonadotropin transcript and sex steroid levels is consistent with the higher gonadotrope cell numbers seen in the adult pituitaries. This indicates that the increase in gonadotropin production is not only via increased cell activity, but also via increased cell number. Our results agree with our previous studies in medaka in which we reported higher numbers of both Fsh (Fontaine et al., 2020a) and Lh cells (Fontaine et al., 2019) in adults compared to juveniles. They further agree with a study in juvenile male African catfish in which Lh cell number increases with the progression of spermatogenesis (Schulz et al., 1997; Cavaco et al., 2001).

To investigate whether sex steroids actually regulate gonadotrope cell activity and numbers, we optimized a protocol which enables gonadal sex steroid removal and the recovery of its physiological levels. Considering the short half-life of sex steroids (Kayo et al., 2020) and taking into account the observed circadian rhythmicity of circulating sex steroids in medaka, the fish were fed five hours before and sampled two hours after the onset of the dark phase. We found that these two time points were optimal to match the levels in GDX fish with those of control fish at the time of their natural peak. Sex steroid supplemented via water exposure or intraperitoneal implantation could cause extremely high circulating sex steroid levels exceeding physiological levels, or even bioaccumulation (Maunder et al., 2007; Kayo et al., 2020), which might result in different physiological effects. For instance, in medaka, E2 administration via feeding (Kanda et al., 2011) upregulates *lhb* mRNA levels while bathing (Fontaine et al., 2019) downregulates them. Thus, using a protocol of sex steroid administration that restores physiological levels is essential to assess the true effect of sex steroids on animal physiology.

Using an optimized protocol with sex steroid supplemented feed, we found that sex steroids suppress *fshb* levels, which is consistent with a previous study in medaka (Kanda et al., 2011; Kayo et al., 2019) and zebrafish (Golan et al., 2014). However in the present study, sex steroids do not to rescue the decreased *lhb* levels induced by GDX which differs from a previous study (Kanda et al., 2011) in which both E2 and 11-KT rescued *lhb* in adult female medaka. Since neither 11-KT nor E2 rescued *lhb* levels in GDX males and females, respectively, we thus tried a combination of *in vivo* intraperitoneal injection of Gnrh1 and E2 feeding in female medaka. Indeed, co-treatment with Gnrh1 and E2 was found to upregulate *lhb* levels *ex vivo* isolated pituitaries from adult female medaka (Karigo et al., 2012). However, we did not observe any effect on *lhb* transcript levels. Finally, estrogenic effects can be involved in the regulation of *lhb* in males. T can be transformed into E2 by the enzyme aromatase which is highly expressed in the brain and pituitary in fish (for review (Fontaine et al., 2020c)). As T treatment was found to elevate Lh-containing granules in male juvenile African catfish (Cavaco et al., 2001), we fed with T but found that T downregulates *lhb* levels in adult male medaka. Whether this is due to species or stage differences remains to be investigated. The failure of sex steroid supplemented food to restore transcript levels occurred not only for *lhb*, but also for other genes, including aromatase (*cyp19a1b*) and sex steroid receptor genes. Activin-follistatin, which are also produced in the gonads, are suggested to regulate *lhb* levels during spawning period in goldfish (Cheng et al., 2007). Therefore, removal of activin-follistatin and other gonadal factors could explain why certain mRNA levels, such as *lhb*, cannot be recovered just with sex steroid supplementation. On the other hand, it is possible that the second peak of E2 observed during daytime in females (in agreement with (Kayo et al., 2020)) might be needed for the regulation of these genes. While this would provide an explanation for females, but not for males in which only one diurnal peak of 11-KT is observed, this therefore requires further investigation.

Interestingly, not only does the activity of gonadotrope cells change following gonadectomy (GDX) but also population volume. The lack of differences in individual cell volume between groups indicates that the increased Fsh cell population volume in GDX fish results from cell hyperplasia. In both sexes, the number of Fsh cells increases in GDX fish, while the number of Lh cell decreases in females, but not in males. To our knowledge, this is the first study in teleost fish showing evidence of gonadotrope cell hyperplasia as a result of an abrupt change in circulating sex steroid levels. In mammals, gonadotrope cell hyperplasia has been shown in GDX rats (Inoue and Kurosumi, 1981; Ibrahim et al., 1986; Sakai et al., 1988) and mouse (Abel et al., 2013; Durán-Pastén et al., 2013). This suggests that the phenomenon behind the effect of gonadal sex steroid removal on gonadotrope cell hyperplasia is conserved across vertebrate species.

After we found that sex steroids indeed play a role in regulating gonadotrope cell activity and number, we then investigated the origin of the newborn Fsh cells. In mammals and fish, increasing numbers of pituitary endocrine cells are usually associated with one or more of three distinct phenomena: transdifferentiation, cell mitosis, and progenitor cell differentiation (Fontaine et al., 2020b; Fontaine et al., 2020c).

In our previous study using single-cell RNA sequencing, we found cells co-expressing *fshb*/*lhb* and *tshba/fshb* (Royan et al., 2021) that we suggested to be in intermediate stages of transdifferentiation when two sets of endocrine cell markers are expressed (Fontaine et al., 2022). As the stable levels of *caspase3* (an apoptotic marker (Julien and Wells, 2017)) suggest that the reduced number of Lh cells in the female pituitary is not a result of apoptosis, we tested whether Lh cells transdifferentiate to the Fsh phenotype. Indeed, a previous study in medaka demonstrated that Fsh cells started to produce *lhb* when stimulated with Gnrh1 *in vitro* (Fontaine et al., 2020a). However, we did not observe differences in *fshb*/*lhb*-expressing cell numbers between control and GDX fish, suggesting that gonadal sex steroids do not regulate this process. Instead of transdifferentiating, Lh cells might become quiescent due to the absence of sex steroids. A previous study reported elevated Lh-containing granules after T treatment despite no increased cell mitosis (Cavaco et al., 2001), suggesting activation of dormant Lh cells.

Interestingly, removal of gonadal sex steroids drastically increases the number of pituitary cells co-expressing *fshb* and *tshba* in females, but not in males, and this phenotype persists for at least six weeks after GDX. In addition, no change is observed in Tsh cell population volume. These observations indicate that Tsh cells remain bi-hormonal, producing both *tshba* and *fshb*. The *fshb*/*tshba* phenotype is abolished with E2 supplementation, suggesting that gonadal sex steroids represent an important regulator of thyrogonadotropes. The presence of thyrogonadotropes agrees with our previous study showing several *fshb*/*tshba*-expressing cells in adult medaka pituitary (Royan et al., 2021), while the current finding suggests that the removal of the negative sex steroid feedback on Fsh cells drives a phenotypic change (transdifferentiation) of Tsh to Tsh/Fsh to increase Fsh hormone production in females. Fsh is essential for female folliculogenesis, but not for male spermatogenesis, as shown by an *fshb* knock-out study in medaka (Takahashi et al., 2016). Fsh cells juxtapose with Tsh cells that constitute a large population in the adult female pituitary (Royan et al., 2021). This proximity might allow an efficient cell-cell communication between Tsh and Fsh cells, resulting in Tsh cells participating in Fsh production in females. This would allow the pituitary to meet the high demand for Fsh required for folliculogenesis.

We also show that sex steroids downregulate transcript levels of the mitotic cell marker *mki67* (Takeuchi and Okubo, 2013; Sun and Kaufman, 2018). Using BrdU experiments, we corroborate this finding by showing that sex steroids inhibit Fsh cell mitosis in both sexes, as well as Lh cell mitosis in males. On the other hand, previous studies reported a stimulatory effect of water-borne sex steroid exposure on Fsh-(Fontaine et al., 2020a) and Lh-induced mitosis in medaka (Fontaine et al., 2019), indicating differential regulation depending on administration method. In mammals, gonadotrope cell proliferation has also been observed after GDX (Sakuma et al., 1984; Durán-Pastén et al., 2013), suggesting conserved mechanisms of mitosis among vertebrates. Interestingly, the fact that removal of gonadal sex steroids stimulates Lh cell mitosis in males, but not in females, might explain why the Lh cell number remains constant in males but decreases in females.

Lastly, we evaluated whether progenitor cells, under the control of sex steroids, play a role in gonadotrope cell remodeling. In mammals, it has been shown that *Sox2*-expressing cells can differentiate into all endocrine cell types (Fauquier et al., 2008). While progenitor cells are suggested to temporarily highly proliferative in the pituitary of GDX male rats (Nolan and Levy, 2006), we observed reduced *sox2* levels and Sox2 cell populations in GDX male medaka, suggesting species differences. The SOX2 marker has been found to quickly disappear in mouse progenitor cells during differentiation (Fauquier et al., 2008). Although the reduction of the Sox2 progenitor population observed after GDX suggests they may be recruited to contribute to the Fsh cell hyperplasia, this needs to be confirmed with further experiments, such as cell lineage tracing.

On the other hand, during embryonic development of the mouse pituitary, *cga* expression precedes that of the beta sub-units, *fshb* and *lhb* (Japón et al., 1994; Pope et al., 2006), suggesting its role in gonadotrope cell ontogeny. Although *cga* levels are sex steroid dependent, we did not observe any cell expressing *cga* without *tshb, lhb* or *fshb* in adult medaka. However, whether *cga* is also a differentiating marker in fish is unknown. We thus cannot exclude that new gonadotrope cells are not recruited from progenitors in response to change in circulating sex steroids as suggested by the decrease in Sox2 immunolabeling in males. Furthermore, the higher *cga* expression observed in the Tsh cell population in GDX fish indicates an increasing activity of Tsh cells and their transdifferentiation into Tsh-Fsh cells. Since the Cga subunit is required for both Fsh and Tsh, the *de novo* production of Fsh hormones would necessitate an increase of *cga* expression in the Tsh cells. Therefore, this agrees with the increasing number of cells co-labeled for *fshb* and *tshba* detected with the FISH, suggesting that Tsh cells quickly transdifferentiate following GDX, and retain both Tsh and Fsh hormone production one month after GDX.

Taken together, we come up with a hypothesis as illustrated in Fig. 7. Low levels of sex steroids enable Fsh cells to produce high levels of Fsh, which is necessary for gonadal development. High Fsh production is driven by increasing cell activity (transcript level), and increased cell numbers via cell mitosis in both sexes and transdifferentiation from Tsh cells in females. When the gonad is developed and the gametes mature, the gonad produces high levels of sex steroids that inhibit Fsh cell activity and proliferation. Gonadal sex steroids also regulate Lh cell mitosis in males but not in females, by increasing the number of mitotic Lh cells. Finally, GDX reduces progenitor cell population in males but their role in gonadotrope cell remodeling remains to be determined.

**Figure 7.**
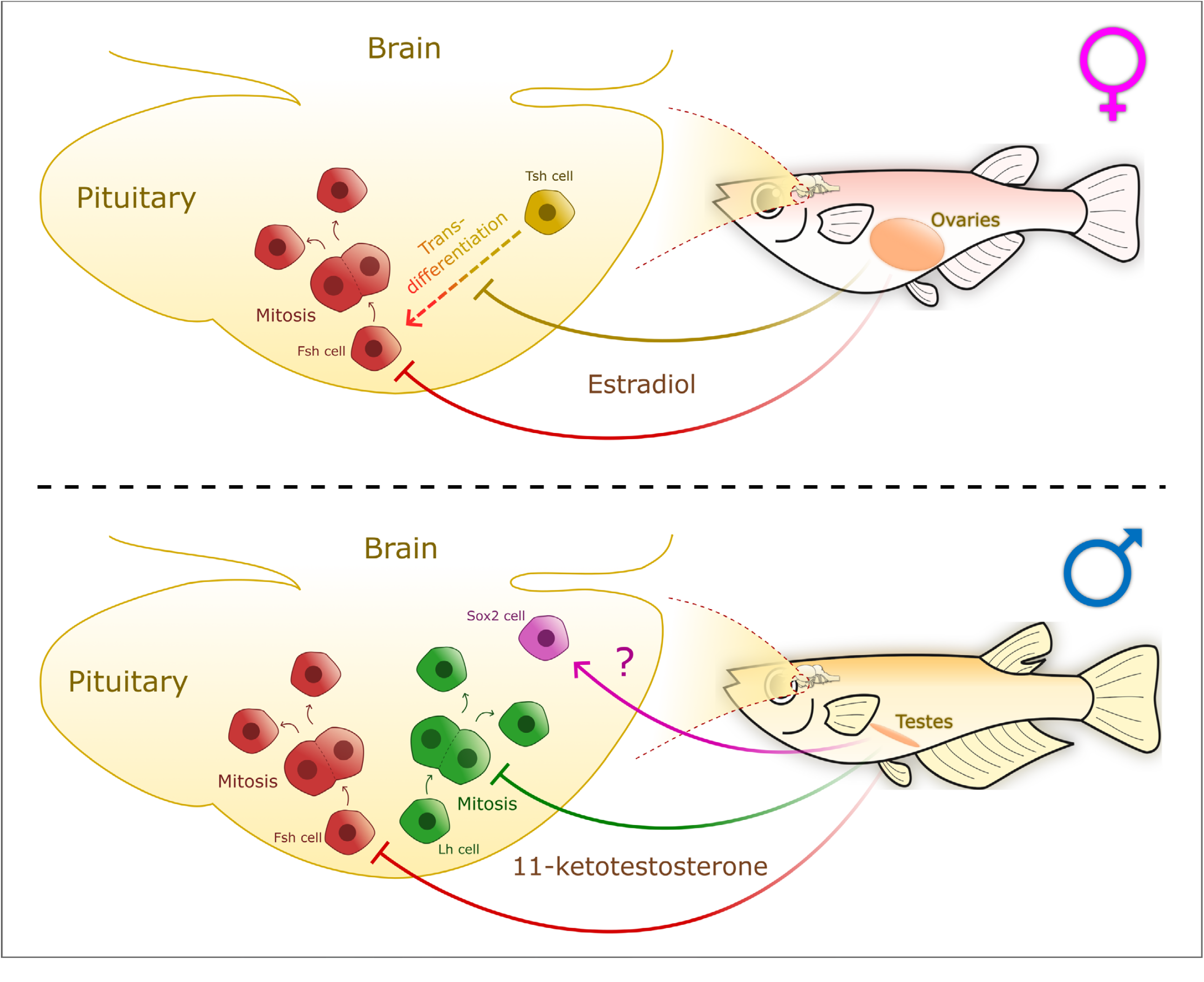
Schematic illustration resuming the sexual dimorphic sex steroid-mediated regulation of gonadotrope cells in medaka. In both sexes, sex steroids inhibit Fsh cell mitosis, while for Lh cells the inhibition only occurs in males. In female, sex steroids inhibit Tsh transdifferentiation to Fsh cells, while in males the absence of sex steroids reduces Sox2 progenitor cell population volume, suggesting that they are recruited to form new cells.

## DECLARATIONS

Ethics approval

Animal experiments were performed according to the recommendations of the care and welfare of research animals at the Norwegian University of Life Sciences.

## COMPETING INTERESTS

The authors declare to have no competing financial interests.

## FUNDING

This work was funded by the Norwegian University of Life Sciences to RF, and Japan Society for the Promotion of Science (JSPS) Grant 20K22587 to DK.

## AUTHOR’S CONTRIBUTIONS

MRR carried out all experimental work. All authors participated in the study design, and the analysis of the results. MRR and RF wrote the manuscript with the input from all other authors.

## ACKNOWLEDGEMENTS

We thank Dr. Shinji Kanda for his inputs on the protocol optimization. We are grateful to Dr. Tomoya Nakayama who kindly provided us the *cga* plasmid. We also thank Anthony Peltier, Lourdes Carreon G Tan, and Arturas Kavaliauskis for fish facility maintenance, and Prof. Dianne M. Baker for reviewing and editing the text.

## FIGURE LEGENDS

**Supplementary Figure 1.**
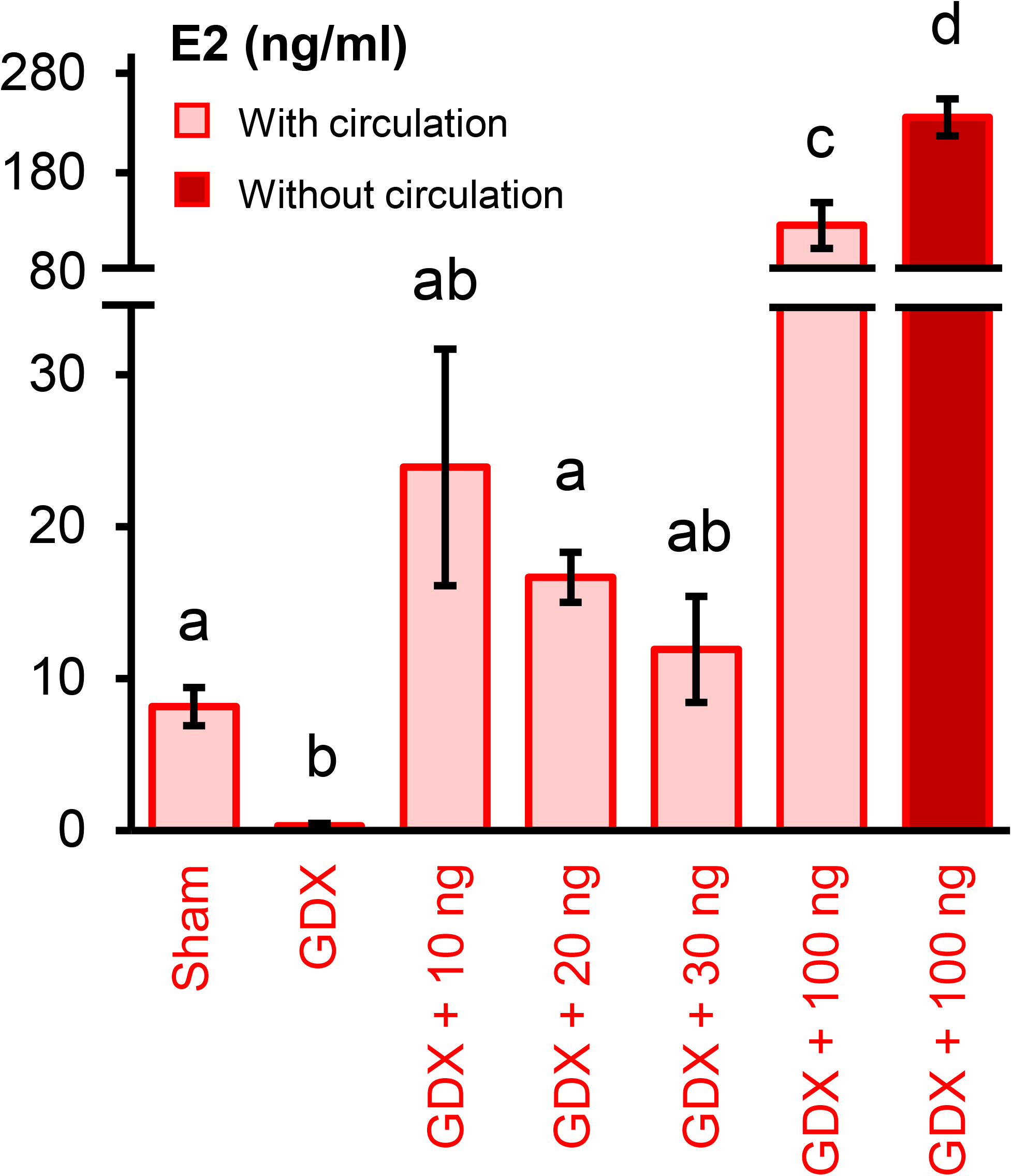
Optimization of sex steroid supplementation protocol. Seven groups of fish belonging to sham-operation, gonadectomized (GDX), GDX + 10 ng/day E2, GDX + 20 ng/day E2, GDX + 30 ng/day E2, and GDX + 100 ng/day E2 with and without circulation were tested to determine the level of E2 that closely mimics the control (sham-operated group) during 5-day experiment (n = 4 - 6). The data were evaluated with One-way ANOVA followed by Tukey *post hoc* test. Graphs are provided as mean ± SEM and different letters display statistical differences (*p* < 0.05) between groups.

**Supplementary Figure 2.**
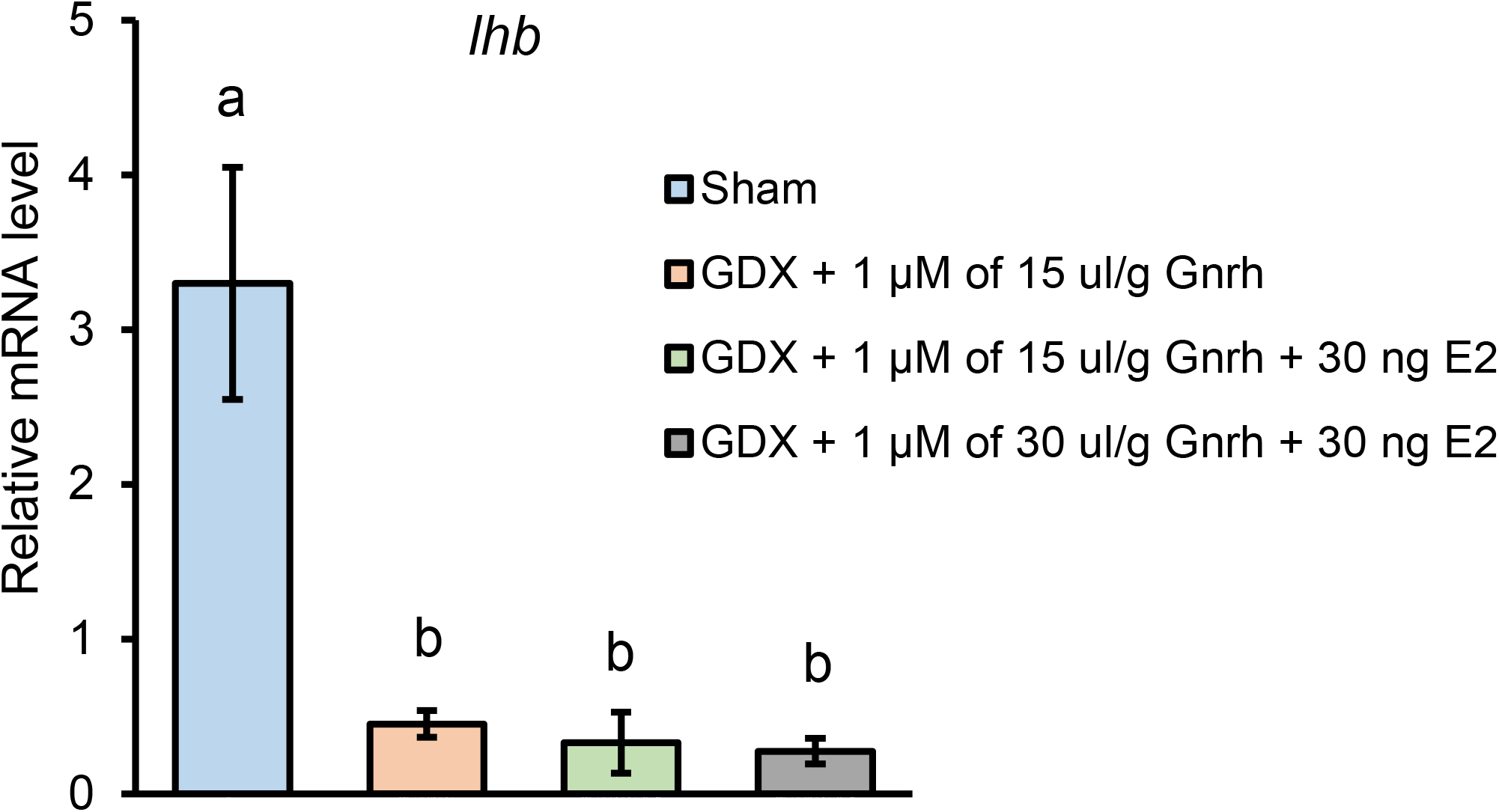
Intraperitoneal injection of Gnrh1 together with E2 supplementation in GDX female medaka does not rescue *lhb* mRNA levels in the pituitary. We injected 1 μM Gnrh1 (Bachem) intraperitoneally as previously performed in zebrafish (Ceriani and Whitlock, 2021). A group of fish is divided into sham-operation, GDX + 15 μl/g of 1 μM Gnrh, GDX + 15 μl/g of 1 μM Gnrh + 30 ng E2, and GDX + 30 μl/g of 1 μM Gnrh + 30 ng E2. Gnrh1 was injected 16 hours (Karigo et al., 2012), while sex steroid feed was administered 7 hours before sampling. Graphs are provided as mean ± SEM, and different letters display statistical differences (*p* < 0.05) between groups as evaluated by One-way ANOVA followed by Tukey *post hoc* test (n = 4).

**Supplementary Figure 3.**
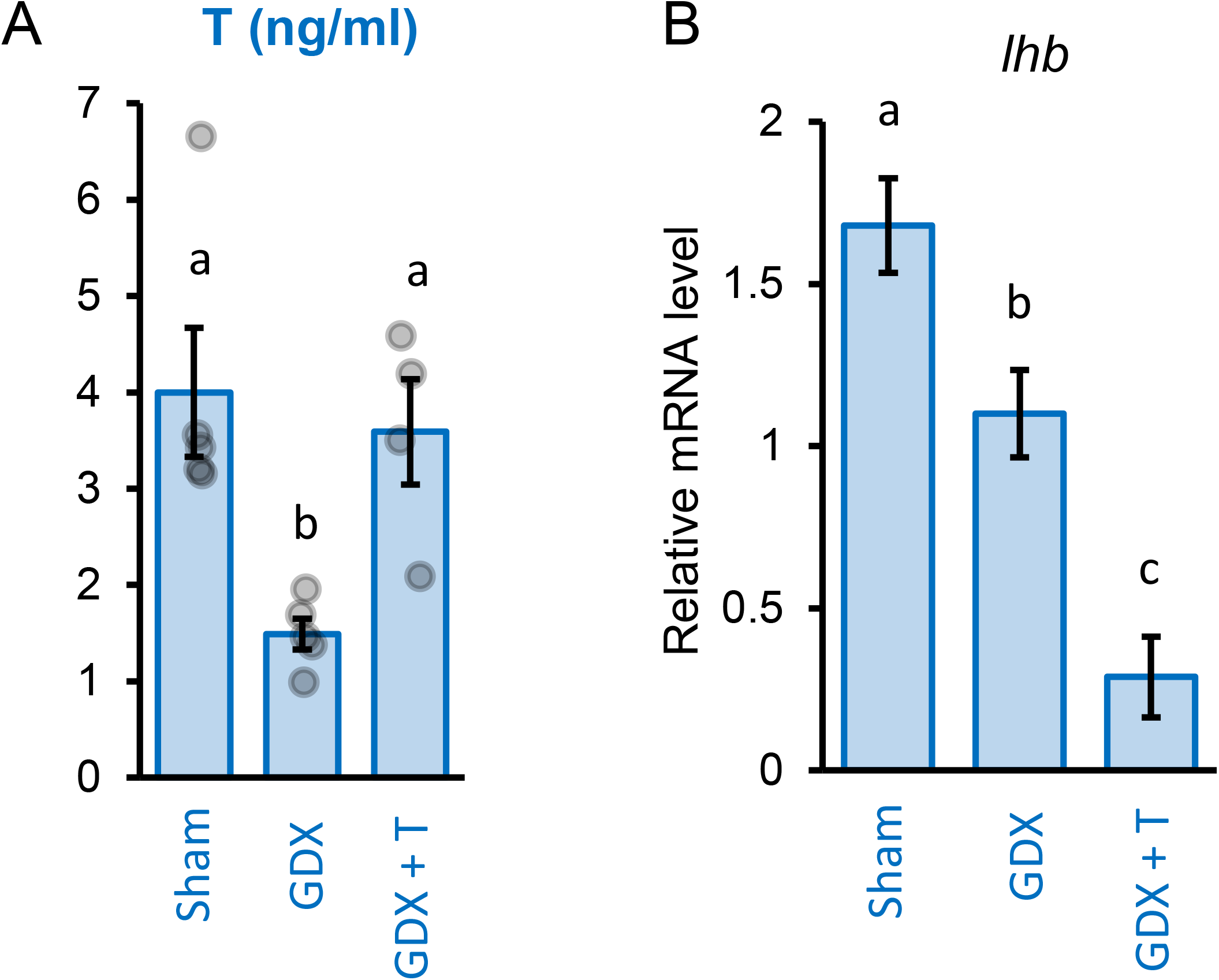
Testosterone (T) treatment does not rescue *lhb* mRNA levels. T supplementation was performed according to the optimized protocol using 30 ng/day of T to the GDX males. **(A)** T levels in sham-operated (Sham), gonadectomized (GDX), and fish fed with T supplemented feed (GDX + T) (n = 4 - 5). **(B)** *lhb* mRNA levels in sham, GDX, and GDX + T (n = 6 - 7). Graphs are provided as mean ± SEM, and different letters display statistical differences (*p* < 0.05) between groups as evaluated by One-way ANOVA followed by Tukey *post hoc* test.

**Supplementary Figure 4.**
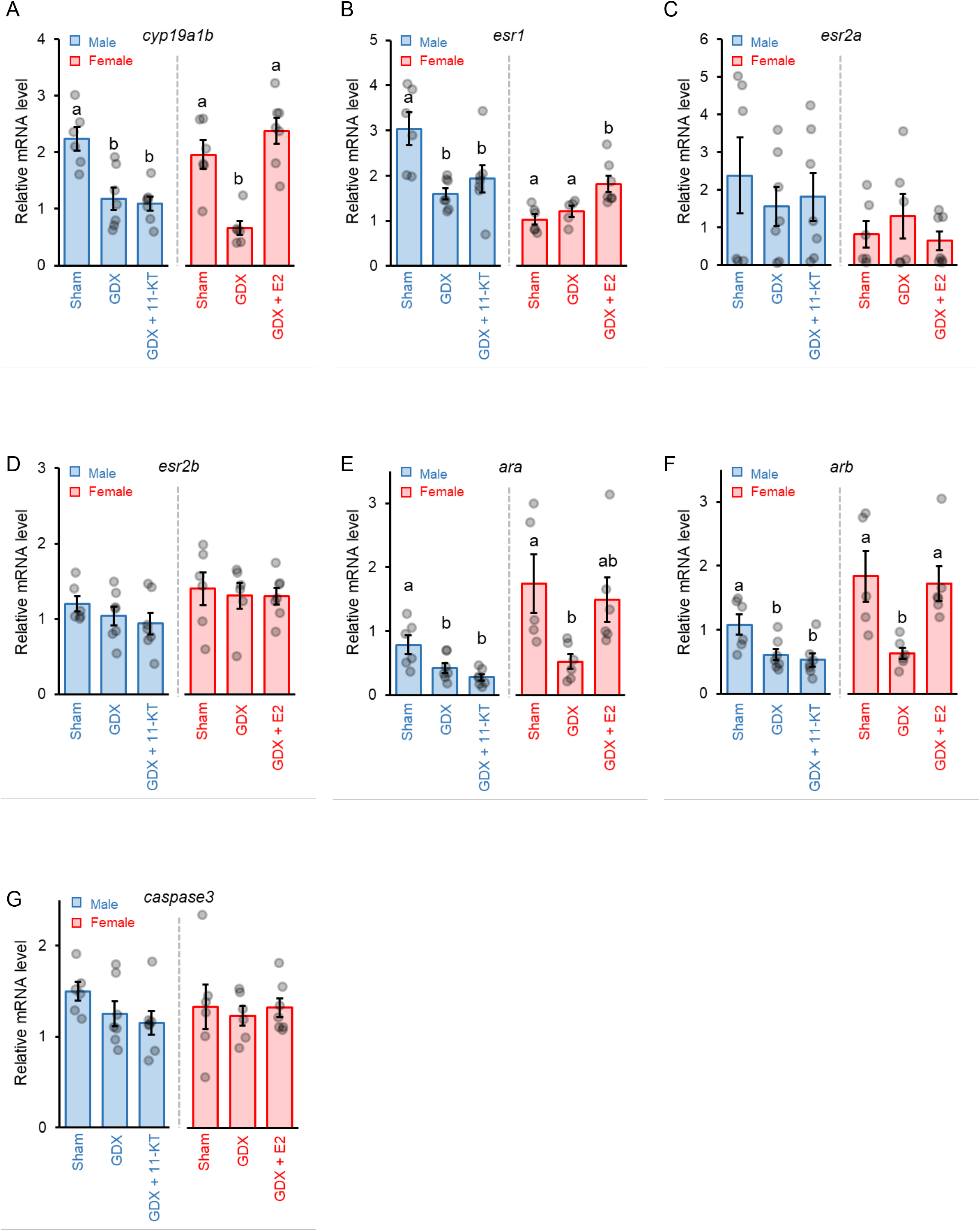
Relative mRNA level of other genes that shows no recovery after sex steroid supplementation. **(A-G)** mRNA levels of *cyp19a1b, esr1, esr2a, esr2b, ara, arb*, and *caspase3* in sham-operated (Sham), gonadectomized (GDX), and fish fed with sex steroids supplemented feed (GDX + 11-KT or GDX + E2). Graphs are provided as mean ± SEM, with jitter dots as the data points. Different letters display statistical differences (*p* < 0.05) between groups as evaluated by One-way ANOVA followed by Tukey *post hoc* test (n = 6 - 7).

**Supplementary Figure 5.**
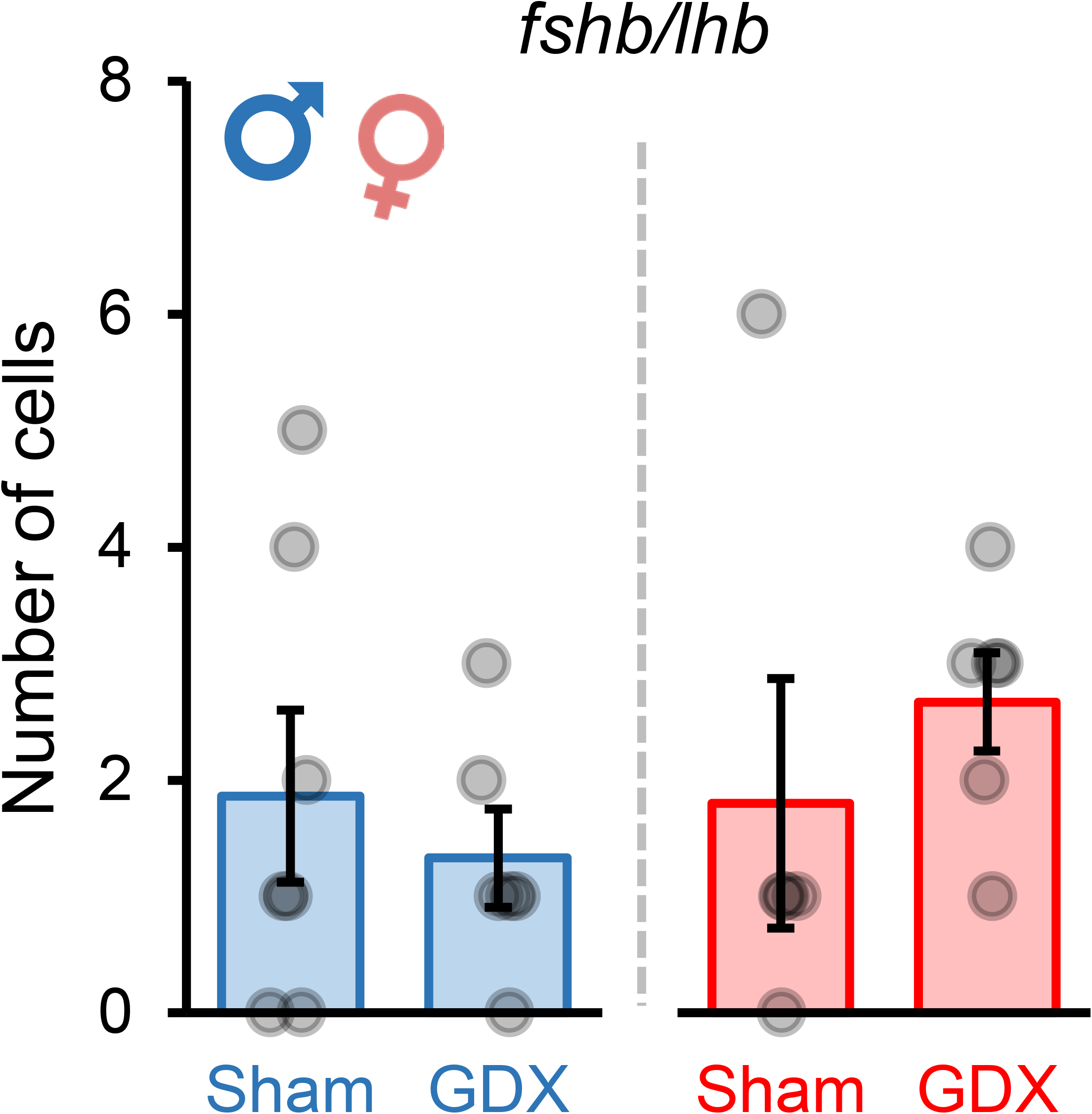
Number of *fshb*/*lhb*-expressing cells in the pituitary of sham and GDX medaka. Number of cells co-expressing *fshb* and *lhb* in sham-operated (Sham) and gonadectomized (GDX) male and female medaka pituitary following fluorescence *in situ* hybridization. Statistically significant differences were evaluated using independent sample student’s t-test (n = 5 - 7). Graphs are provided as mean ± SEM, with jitter dots as the data points.

**Supplementary Figure 6.**
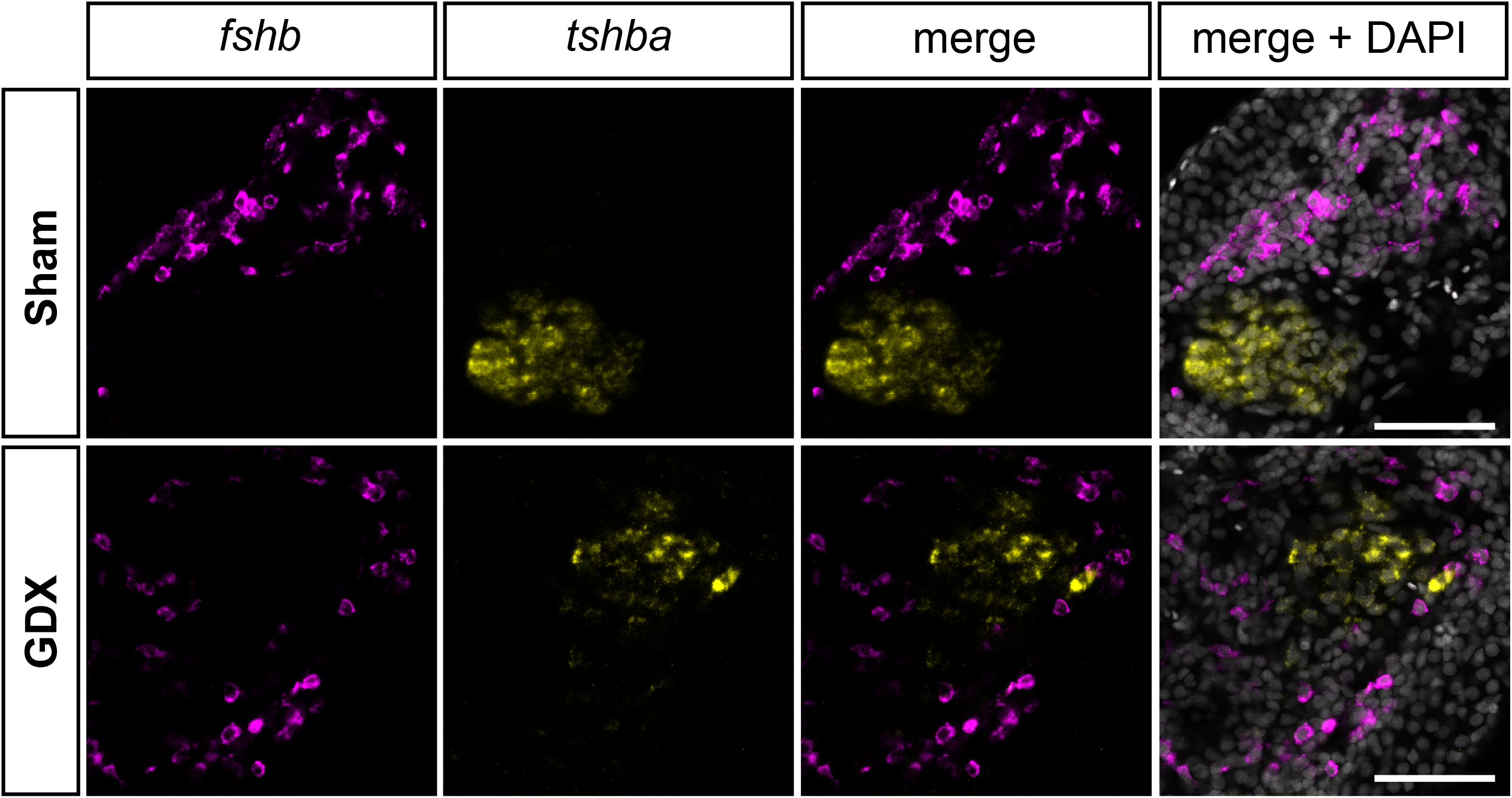
Expression of *fshb* and *tshba* in the male medaka pituitary. Representative confocal planes from five pituitary parasagittal sections labelled for *fshb* (magenta) and *tshba* (yellow) in males. Scale bars are in 50 μm.

**Supplementary Figure 7.**
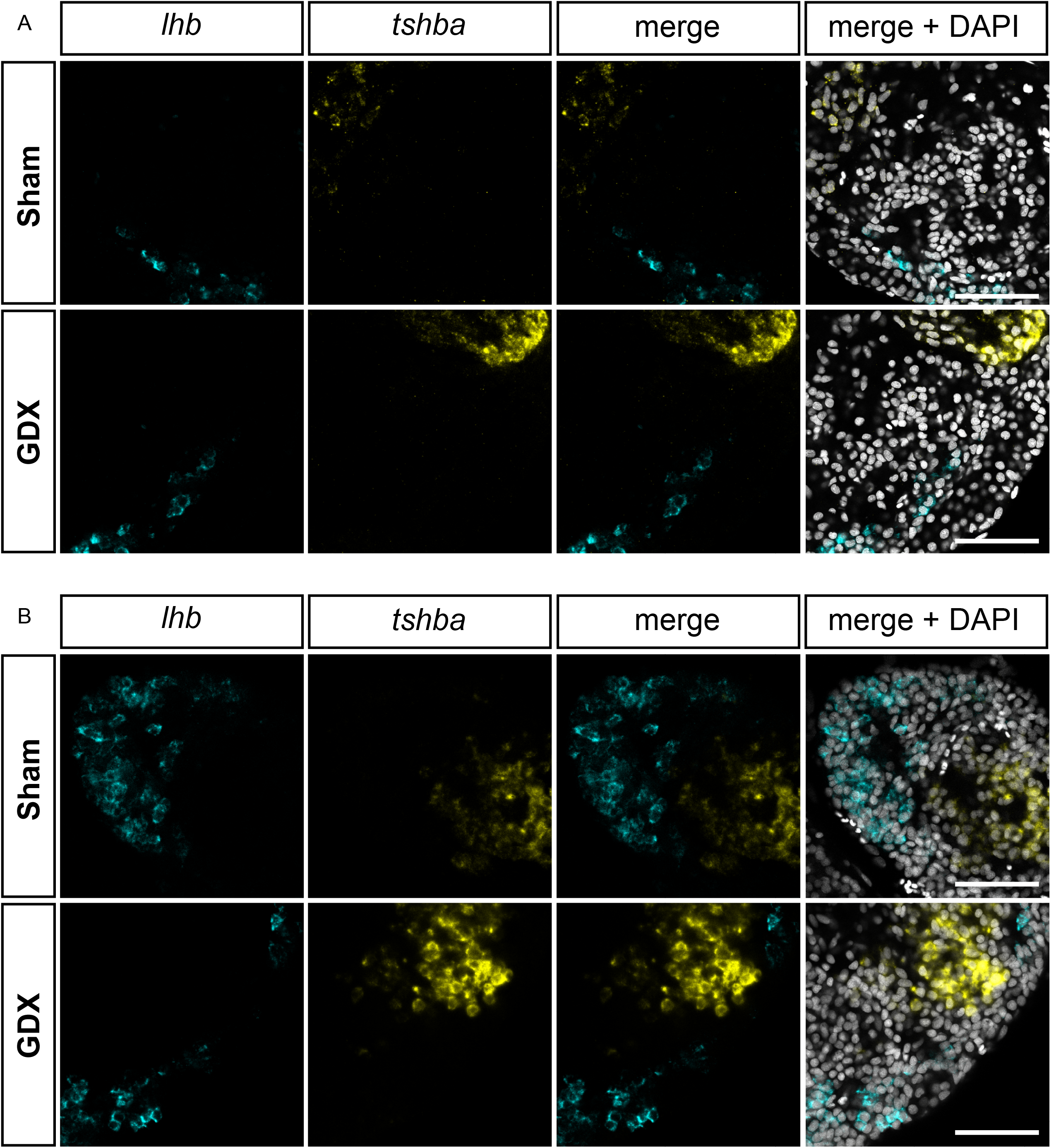
Expression of *lhb* and *tshba* in the medaka pituitary. Representative confocal planes pituitary parasagittal sections labelled for *lhb* (cyan) and *tshba* (yellow) in male **(A)** and female **(B)** medaka (n = 5). Nuclei (grey) were labeled with DAPI staining. Scale bars are in 50 μm.

**Supplementary Figure 8.**
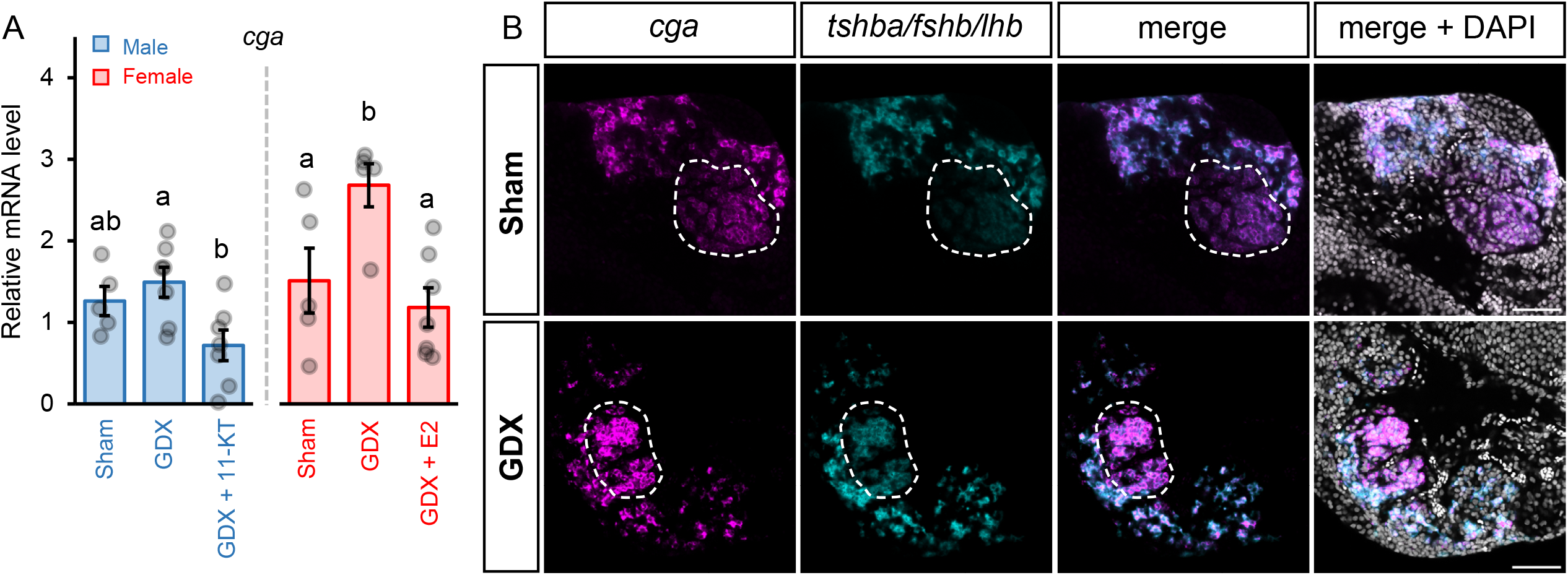
*cga* expression in adult female medaka pituitary. **(A)** *cga* mRNA levels in sham-operated (Sham), gonadectomized (GDX) fish, and fish fed with sex steroids supplemented feed (GDX + 11-KT or GDX + E2) (n = 5 - 7). Statistically significant differences were tested by One-way ANOVA followed by Tukey *post hoc* test. Different letters display statistically significant differences (*p* < 0.05) between groups. **(B)** Representative confocal planes of pituitary parasagittal sections labelled for *cga* (magenta) and combined *tshba*/*fshb*/*lhb* (cyan) in female medaka (n = 5). White dashed line delimits the Tsh cell population (expected according to 3D atlas medaka pituitary (Royan et al., 2021)). Scale bars are in 50 μm.

## Notes

### Competing Interest Statement

The authors have declared no competing interest.

## REFERENCES

Abel, M.H., Charlton, H.M., Huhtaniemi, I., Pakarinen, P., Kumar, T.R., and Christian, H.C. (2013). An Investigation into Pituitary Gonadotrophic Hormone Synthesis, Secretion, Subunit Gene Expression and Cell Structure in Normal and Mutant Male Mice. Journal of Neuroendocrinology 25(10), 863–875. doi: https://doi.org/10.1111/jne.12081.

Burow, S., Fontaine, R., von Krogh, K., Mayer, I., Nourizadeh-Lillabadi, R., Hollander-Cohen, L., et al. (2019). Medaka follicle-stimulating hormone (Fsh) and luteinizing hormone (Lh): Developmental profiles of pituitary protein and gene expression levels. General and Comparative Endocrinology 272, 93–108. doi: https://doi.org/10.1016/j.ygcen.2018.12.006.

Cavaco, J.E.B., van Baal, J., van Dijk, W., Hassing, G.A.M., Th. Goos, H.J., and Schulz, R.W. (2001). Steroid Hormones Stimulate Gonadotrophs in Juvenile Male African Catfish (Clarias gariepinus)1. Biology of Reproduction 64(5), 1358–1365. doi: 10.1095/biolreprod64.5.1358 %J Biology of Reproduction.

Ceriani, R., and Whitlock, K.E. (2021). Gonadotropin Releasing Hormone (GnRH) Triggers Neurogenesis in the Hypothalamus of Adult Zebrafish. International Journal of Molecular Sciences 22(11), 5926.

Chakraborty, T., Shibata, Y., Zhou, L.-Y., Katsu, Y., Iguchi, T., and Nagahama, Y. (2011). Differential expression of three estrogen receptor subtype mRNAs in gonads and liver from embryos to adults of the medaka, Oryzias latipes. Molecular and Cellular Endocrinology 333(1), 47–54. doi: https://doi.org/10.1016/j.mce.2010.12.002.

Cheng, G.F.Y., Yuen, C.-W., and Ge, W. (2007). Evidence for the existence of a local activin–follistatin negative feedback loop in the goldfish pituitary and its regulation by activin and gonadal steroids. Journal of Endocrinology 195(3), 373–384. doi: 10.1677/joe-07-0265.

Devlin, R.H., and Nagahama, Y. (2002). Sex determination and sex differentiation in fish: an overview of genetic, physiological, and environmental influences. Aquaculture 208(3), 191–364. doi: https://doi.org/10.1016/S0044-8486(02)00057-1.

Durán-Pastén, M.L., Fiordelisio-Coll, T., and Hernández-Cruz, A. (2013). Castration-Induced Modifications of GnRH-Elicited [Ca2+]i Signaling Patterns in Male Mouse Pituitary Gonadotrophs In Situ: Studies in the Acute Pituitary Slice Preparation1. Biology of Reproduction 88(2). doi: 10.1095/biolreprod.112.103812.

Fauquier, T., Rizzoti, K., Dattani, M., Lovell-Badge, R., and Robinson, I.C.A.F. (2008). SOX2-expressing progenitor cells generate all of the major cell types in the adult mouse pituitary gland. Proceedings of the National Academy of Sciences 105(8), 2907. doi: 10.1073/pnas.0707886105.

Fontaine, R., Affaticati, P., Yamamoto, K., Jolly, C., Bureau, C., Baloche, S., et al. (2013). Dopamine Inhibits Reproduction in Female Zebrafish (Danio rerio) via Three Pituitary D2 Receptor Subtypes. Endocrinology 154(2), 807–818. doi: https://doi.org/10.1210/en.2012-1759.

Fontaine, R., Ager-Wick, E., Hodne, K., and Weltzien, F.-A. (2019). Plasticity of Lh cells caused by cell proliferation and recruitment of existing cells. Journal of Endocrinology 240(2), 361. doi: https://doi.org/10.1530/JOE-18-0412.

Fontaine, R., Ager-Wick, E., Hodne, K., and Weltzien, F.-A. (2020a). Plasticity in medaka gonadotropes via cell proliferation and phenotypic conversion. Journal of Endocrinology 245(1), 21. doi: https://doi.org/10.1530/JOE-19-0405.

Fontaine, R., Ciani, E., Haug, T.M., Hodne, K., Ager-Wick, E., Baker, D.M., et al. (2020b). Gonadotrope plasticity at cellular, population and structural levels: A comparison between fishes and mammals. General and Comparative Endocrinology 287, 113344. doi: https://doi.org/10.1016/j.ygcen.2019.113344.

Fontaine, R., Rahmad Royan, M., Henkel, C., Hodne, K., Ager-Wick, E., and Weltzien, F.-A. (2022). Pituitary multi-hormone cells in mammals and fish: history, origin, and roles. Frontiers in Neuroendocrinology 67, 101018. doi: https://doi.org/10.1016/j.yfrne.2022.101018.

Fontaine, R., Royan, M.R., von Krogh, K., Weltzien, F.-A., and Baker, D.M. (2020c). Direct and Indirect Effects of Sex Steroids on Gonadotrope Cell Plasticity in the Teleost Fish Pituitary. Frontiers in Endocrinology 11(858). doi: https://doi.org/10.3389/fendo.2020.605068.

Golan, M., Biran, J., and Levavi-Sivan, B. (2014). A Novel Model for Development, Organization, and Function of Gonadotropes in Fish Pituitary. 5(182). doi: https://doi.org/10.3389/fendo.2014.00182.

Hildahl, J., Sandvik, G.K., Lifjeld, R., Hodne, K., Nagahama, Y., Haug, T.M., et al. (2012). Developmental tracing of luteinizing hormone β-subunit gene expression using green fluorescent protein transgenic medaka (Oryzias latipes) reveals a putative novel developmental function. 241(11), 1665–1677. doi: 10.1002/dvdy.23860.

Hodne, K., Fontaine, R., Ager-Wick, E., and Weltzien, F.-A. (2019). Gnrh1-Induced Responses Are Indirect in Female Medaka Fsh Cells, Generated Through Cellular Networks. Endocrinology 160(12), 3018–3032. doi: 10.1210/en.2019-00595 %J Endocrinology.

Hori, H. (2011). “A Glance at the Past of Medaka Fish Biology,” in Medaka: A Model for Organogenesis, Human Disease, and Evolution, eds. K. Naruse, M. Tanaka & H. Takeda. (Tokyo: Springer Japan), 1–16.

Ibrahim, S.N., Moussa, S.M., and Childs, G.V. (1986). Morphometric Studies of Rat Anterior Pituitary Cells after Gonadectomy: Correlation of Changes in Gonadotropes with the Serum Levels of Gonadotropins*. Endocrinology 119(2), 629–637. doi: 10.1210/endo-119-2-629.

Inoue, K., and Kurosumi, K. (1981). Mode of Proliferation of Gonadotrophic Cells of the Anterior Pituitary after Castration-Immunocytochemical and Autoradiographic Studies. Archivum histologicum japonicum 44(1), 71–85. doi: 10.1679/aohc1950.44.71.

Jamovi (2021). “jamovi (Version 2.2.5)”, in: The Jamovi Project. (Sydney, Australia: The Jamovi Project).

Japón, M.A., Rubinstein, M., and Low, M.J. (1994). In situ hybridization analysis of anterior pituitary hormone gene expression during fetal mouse development. Journal of Histochemistry & Cytochemistry 42(8), 1117–1125. doi: 10.1177/42.8.8027530.

Julien, O., and Wells, J.A. (2017). Caspases and their substrates. Cell Death & Differentiation 24(8), 1380–1389. doi: 10.1038/cdd.2017.44.

Kanda, S. (2019). Evolution of the regulatory mechanisms for the hypothalamic-pituitary-gonadal axis in vertebrates–hypothesis from a comparative view. General and Comparative Endocrinology 284, 113075. doi: https://doi.org/10.1016/j.ygcen.2018.11.014.

Kanda, S., Akazome, Y., Matsunaga, T., Yamamoto, N., Yamada, S., Tsukamura, H., et al. (2008). Identification of KiSS-1 Product Kisspeptin and Steroid-Sensitive Sexually Dimorphic Kisspeptin Neurons in Medaka (Oryzias latipes). Endocrinology 149(5), 2467–2476. doi: https://doi.org/10.1210/en.2007-1503.

Kanda, S., Okubo, K., and Oka, Y. (2011). Differential regulation of the luteinizing hormone genes in teleosts and tetrapods due to their distinct genomic environments – Insights into gonadotropin beta subunit evolution. General and Comparative Endocrinology 173(2), 253–258. doi: https://doi.org/10.1016/j.ygcen.2011.05.015.

Karigo, T., Kanda, S., Takahashi, A., Abe, H., Okubo, K., and Oka, Y. (2012). Time-of-Day-Dependent Changes in GnRH1 Neuronal Activities and Gonadotropin mRNA Expression in a Daily Spawning Fish, Medaka. Endocrinology 153(7), 3394–3404. doi: 10.1210/en.2011-2022.

Kawabata-Sakata, Y., Nishiike, Y., Fleming, T., Kikuchi, Y., and Okubo, K. (2020). Androgen-dependent sexual dimorphism in pituitary tryptophan hydroxylase expression: relevance to sex differences in pituitary hormones. Proceedings of the Royal Society B: Biological Sciences 287(1928), 20200713. doi: https://doi.org/10.1098/rspb.2020.0713.

Kayo, D., Oka, Y., and Kanda, S. (2020). Examination of methods for manipulating serum 17β-Estradiol (E2) levels by analysis of blood E2 concentration in medaka (Oryzias latipes). General and Comparative Endocrinology 285, 113272. doi: https://doi.org/10.1016/j.ygcen.2019.113272.

Kayo, D., Zempo, B., Tomihara, S., Oka, Y., and Kanda, S. (2019). Gene knockout analysis reveals essentiality of estrogen receptor β1 (Esr2a) for female reproduction in medaka. Scientific Reports 9(1), 8868. doi: https://doi.org/10.1038/s41598-019-45373-y.

Kazeto, Y., and Trant, J.M. (2005). Molecular biology of channel catfish brain cytochrome P450 aromatase (CYP19A2): cloning, preovulatory induction of gene expression, hormonal gene regulation and analysis of promoter region. Journal of Molecular Endocrinology 35(3), 571–583. doi: 10.1677/jme.1.01805.

Kinoshita, M., Murata, K., Naruse, K., and Tanaka, M. (2009). “Looking at Adult Medaka,” in Medaka: Biology, Management, and Experimental Protocols, eds. M. Kinoshita, K. Murata, K. Naruse & M. Tanaka. Wiley Blackwell), 117–164.

Klenke, U., and Zohar, Y. (2003). Gonadal regulation of gonadotropin subunit expression and pituitary Lh protein content in female hybrid striped bass. Fish Physiology and Biochemistry 28(1), 25–27. doi: 10.1023/B:FISH.0000030465.56876.97.

Matsuda, M., Nagahama, Y., Shinomiya, A., Sato, T., Matsuda, C., Kobayashi, T., et al. (2002). DMY is a Y-specific DM-domain gene required for male development in the medaka fish. Nature 417(6888), 559–563. doi: 10.1038/nature751.

Maunder, R.J., Matthiessen, P., Sumpter, J.P., and Pottinger, T.G. (2007). Rapid bioconcentration of steroids in the plasma of three-spined stickleback Gasterosteus aculeatus exposed to waterborne testosterone and 17β-oestradiol. Journal of Fish Biology 70(3), 678–690. doi: https://doi.org/10.1111/j.1095-8649.2006.01303.x.

Nakane, P.K. (1970). Classifications of anterior pituitary cell types with immunoenzyme histochemistry. Journal of Histochemistry & Cytochemistry 18(1), 9–20. doi: https://doi.org/10.1177/18.1.9.

Nanda, I., Kondo, M., Hornung, U., Asakawa, S., Winkler, C., Shimizu, A., et al. (2002). A duplicated copy of DMRT1 in the sex-determining region of the Y chromosome of the medaka, Oryzias latipes. Proceedings of the National Academy of Sciences 99(18), 11778–11783. doi: doi:10.1073/pnas.182314699.

Nolan, L.A., and Levy, A. (2006). A Population of Non-Luteinising Hormone/Non-Adrenocorticotrophic Hormone-Positive Cells in the Male Rat Anterior Pituitary Responds Mitotically to Both Gonadectomy and Adrenalectomy. Journal of Neuroendocrinology 18(9), 655–661. doi: https://doi.org/10.1111/j.1365-2826.2006.01459.x.

Pope, C., McNeilly, J.R., Coutts, S., Millar, M., Anderson, R.A., and McNeilly, A.S. (2006). Gonadotrope and thyrotrope development in the human and mouse anterior pituitary gland. Developmental Biology 297(1), 172–181. doi: https://doi.org/10.1016/j.ydbio.2006.05.005.

Powell, J.F.F., Krueckl, S.L., Collins, P.M., and Sherwood, N.M. (1996). Molecular forms of GnRH in three model fishes: rockfish, medaka and zebrafish. 150(1), 17. doi: 10.1677/joe.0.1500017.

Royan, M.R., Kanda, S., Kayo, D., Song, W., Ge, W., Weltzien, F.-A., et al. (2020). Gonadectomy and Blood Sampling Procedures in the Small Size Teleost Model Japanese Medaka (Oryzias latipes). JoVE (166), e62006. doi: https://dx.doi.org/10.3791/62006.

Royan, M.R., Siddique, K., Csucs, G., Puchades, M.A., Nourizadeh-Lillabadi, R., Bjaalie, J.G., et al. (2021). 3D Atlas of the Pituitary Gland of the Model Fish Medaka (Oryzias latipes). Frontiers in Endocrinology 12(912). doi: https://doi.org/10.3389/fendo.2021.719843.

Sakai, T., Inoue, K., Hasegawa, Y., and Kurosumi, K. (1988). Effect of Passive Immunization to Gonadotropin-Releasing Hormone (GnRH) Using GnRH Antiserum on the Mitotic Activity of Gonadotrophs in Castrated Male Rats*. Endocrinology 122(6), 2803–2808. doi: 10.1210/endo-122-6-2803.

Sakuma, S., Shirasawa, N., and Yoshimura, F. (1984). A histometrical study of immunohistochemically identified mitotic adenohypophysial cells in immature and mature castrated rats. Journal of Endocrinology 100(3), 323–328. doi: 10.1677/joe.0.1000323.

Saligaut, C., Linard, B., Mañanos, E.L., Kah, O., Breton, B., and Govoroun, M. (1998). Release of Pituitary Gonadotrophins GtH I and GtH II in the Rainbow Trout (Oncorhynchus mykiss): Modulation by Estradiol and Catecholamines. General and Comparative Endocrinology 109(3), 302–309. doi: https://doi.org/10.1006/gcen.1997.7033.

Schulz, R.W., Zandbergen, M.A., Peute, J., Bogerd, J., van Dijk, W., and Goos, H.J.T. (1997). Pituitary Gonadotrophs are Strongly Activated at the Beginning of Spermatogenesis in African Catfish, Clarias Gariepinus. Biology of Reproduction 57(1), 139–147. doi: 10.1095/biolreprod57.1.139.

Sohn, Y.C., Suetake, H., Yoshiura, Y., Kobayashi, M., and Aida, K. (1998). Structural and expression analyses of gonadotropin Iβ subunit genes in goldfish (Carassius auratus). Gene 222(2), 257–267. doi: https://doi.org/10.1016/S0378-1119(98)00505-8.

Sun, X., and Kaufman, P.D. (2018). Ki-67: more than a proliferation marker. Chromosoma 127(2), 175–186. doi: 10.1007/s00412-018-0659-8.

Takahashi, A., Kanda, S., Abe, T., and Oka, Y. (2016). Evolution of the Hypothalamic-Pituitary-Gonadal Axis Regulation in Vertebrates Revealed by Knockout Medaka. Endocrinology 157(10), 3994–4002. doi: 10.1210/en.2016-1356.

Takeuchi, A., and Okubo, K. (2013). Post-Proliferative Immature Radial Glial Cells Female-Specifically Express Aromatase in the Medaka Optic Tectum. PLOS ONE 8(9), e73663. doi: 10.1371/journal.pone.0073663.

Vetillard, A., Atteke, C., Saligaut, C., Jego, P., and Bailhache, T. (2003). Differential regulation of tyrosine hydroxylase and estradiol receptor expression in the rainbow trout brain. Molecular and Cellular Endocrinology 199(1), 37–47. doi: https://doi.org/10.1016/S0303-7207(02)00305-2.

Weltzien, F.-A., Andersson, E., Andersen, Ø., Shalchian-Tabrizi, K., and Norberg, B. (2004). The brain– pituitary–gonad axis in male teleosts, with special emphasis on flatfish (Pleuronectiformes). Comparative Biochemistry and Physiology Part A: Molecular & Integrative Physiology 137(3), 447–477. doi: https://doi.org/10.1016/j.cbpb.2003.11.007.

Weltzien, F.-A., Hildahl, J., Hodne, K., Okubo, K., and Haug, T.M. (2014). Embryonic development of gonadotrope cells and gonadotropic hormones – Lessons from model fish. Molecular and Cellular Endocrinology 385(1), 18–27. doi: https://doi.org/10.1016/j.mce.2013.10.016.

Wittbrodt, J., Shima, A., and Schartl, M. (2002). Medaka — a model organism from the far east. Nature Reviews Genetics 3(1), 53–64. doi: https://doi.org/10.1038/nrg704.

Yaron, Z., and Levavi-Sivan, B. (2011). “Hormonal control of reproduction and growth | Endocrine Regulation of Fish Reproduction,” in Encyclopedia of Fish Physiology, ed. A.P. Farrell. (kSan Diego: Academic Press), 1500–1508.

